# Eukaryotic biodiversity and ecological networks from the surface to the mesopelagic in the Northwest Atlantic Slope Water

**DOI:** 10.1101/2025.05.12.653512

**Authors:** Nina Yang, Elizabeth A. Allan, Sarah E. Stover, Benjamin D. Grassian, Heidi M. Sosik, Joel K. Llopiz, Annette F. Govindarajan

## Abstract

The diversity and interactions among mesopelagic organisms are difficult to study and as a result, are insufficiently unaccounted for in food web and biogeochemical models. This knowledge gap hinders our ability to model and forecast ecosystem function and formulate effective policies for conservation and management in the face of growing interest in exploiting midwater living resources. We used multi-marker metabarcoding of environmental DNA (eDNA) samples collected from Northwest Atlantic Slope Water to resolve patterns of eukaryotic community composition spanning taxonomically across protists (microbial eukaryotes), invertebrates, and vertebrates and vertically from the ocean surface to the base of the mesopelagic zone. With statistical network analyses, we explored cross-kingdom associations including putative trophic dynamics such as food web interactions and evaluated network robustness to biodiversity loss. We found depth-specific communities of distinct protist, invertebrate, and vertebrate assemblages. Ecological networks for the epipelagic, upper mesopelagic, and lower mesopelagic suggest that protists are keystone taxa and important mediators of trophic interactions; they increase network complexity and contribute to network stability. We also identified metazoans including copepods, gelatinous taxa (cnidarians, tunicates), and mesopelagic fish as important components of network interactions. Our study demonstrates a holistic approach to generate insights on mesopelagic biodiversity and implications for ecosystem resilience that can inform ocean governance.

## 2. INTRODUCTION

In the vast mesopelagic region of the ocean from the base of the sunlit zone to 1000 m deep (Buesseler et al., 2020), diverse ecological communities underpin ecosystem function and critical ecosystem services (Oliver et al., 2015; Sala et al., 2021; Nauta and De Domenico, 2024). A wide variety of life, from microscopic phytoplankton to apex predators, comprise complex food webs that support commercially and ecologically important species including sharks, tuna, and billfish (Braun et al., 2023; Iglesias et al., 2023; Willis et al., 2025). In addition, mesopelagic biodiversity play an important role in climate regulation by carrying out biogeochemical processes that determine carbon flux into deeper waters and thus, its long-term sequestration (e.g., 10-1000 years) (Herndl and Reinthaler, 2013; Mazuecos et al., 2015; Cavan et al., 2019; Nguyen et al., 2022).

The mesopelagic zone is intrinsically linked to the surface ocean or epipelagic zone (< 200 m) where phytoplankton harness energy from the sun to convert carbon dioxide into organic carbon. Zooplankton grazing and senescence of primary production can export organic carbon out of the epipelagic, providing a food source for life in the deep ocean (Cawley et al., 2021; Hemsley et al., 2023) and contributing to the largest reservoir of organic carbon on earth (Arístegui et al., 2009). In addition, animals that migrate from the depths to feed at the surface actively transport carbon into deeper waters through diel vertical migration (DVM) behaviors, further demonstrating the interconnected nature of the epipelagic and mesopelagic ocean (Robinson et al., 2010; Davison et al., 2013; Thibault et al., 2025).

Despite the recognized importance of the deep sea, biodiversity and ecological studies have primarily focused on coastal or epipelagic ecosystems, leaving the mesopelagic both undersampled and understudied (Webb et al., 2010; St. John et al., 2016). These knowledge gaps are largely driven by challenges associated with accessing and sampling a vast ecosystem that spans considerable horizontal and vertical spatial scales (Sutton et al., 2017). Difficulty in acquiring logistical resources and expertise for deep sea exploration and research are also major contributing factors (Tolochko and Vadrot, 2021; Amon et al., 2022b). Furthermore, many studies focus on specific groups of organisms in isolation, such as those exclusively examining microbes or metazoans (Yang et al., 2024), which skews our understanding of biodiversity, ecological interactions, and functional roles.

Growing interest in mesopelagic fisheries for aquaculture and nutraceuticals (Martin et al., 2020; Bisson et al., 2023), deep sea mining activities (Drazen et al., 2020; Amon et al., 2022a; Dowd et al., 2025), and marine carbon dioxide removal efforts (Hernández-León, 2023; Oschlies et al., 2025) presents increasing pressures on the mesopelagic and its ecosystem services, including overexploitation and habitat destruction. In the face of these anthropogenic pressures, limited data present significant challenges for effective ocean governance of deep pelagic ecosystems, including conservation, resource management, and environmental impact assessments.

In the last decade, environmental DNA (eDNA) approaches have emerged as a reliable and non-invasive solution to survey biodiversity (McClenaghan et al., 2020; Chavez et al., 2021; Holman et al., 2021; Yeh and Fuhrman, 2022; Govindarajan et al., 2023a; Dan et al., 2024). With eDNA, organisms across all domains of life can be detected from a single water sample by targeting multiple marker genes (Amaral-Zettler et al., 2009; Miya et al., 2015), making eDNA approaches ideal for large-scale biomonitoring across varied spatiotemporal scales (Deiner et al., 2017; Gold et al., 2021, 2022; Blancher et al., 2022; De Brauwer et al., 2023; Thomsen et al., 2024). However, few multi-marker metabarcoding studies have been conducted in deep pelagic ecosystems and existing studies tend to emphasize organismal diversity and distribution rather than interactions among taxa (Giner et al., 2020; Laroche et al., 2020a; Govindarajan et al., 2021; Holman et al., 2021; Govindarajan et al., 2023b). A more holistic approach would facilitate our ability to effectively understand how community composition and organismal interactions across eukaryotic taxa influence crucial ecosystem processes. For example, recent studies have employed multi-marker metabarcoding approaches to resolve diversity patterns from prokaryotes to top predators (Djurhuus et al., 2020; Holman et al., 2021). Using statistical and network analyses, these studies also documented key predator-prey interactions, identified potential keystone taxa, and uncovered unexplored trophic connections. Collectively, these insights provide necessary context for ecosystem health and may serve as important indicators for ecosystem function in a changing ocean.

We applied an eDNA metabarcoding approach with two gene markers to investigate patterns of eukaryotic biodiversity and their interactions to better understand ecosystem structure of the epipelagic and mesopelagic zones in the Northwest Atlantic Slope Water region. We calculated diversity indices to determine depth-specific biodiversity patterns. We then performed covariance network analyses on multi-marker metabarcoding data, constructing depth-specific networks for metazoans (invertebrates and vertebrates), protists, and all taxa combined across the epipelagic (10 m, 100 m), upper mesopelagic (300 m, 500 m), and lower mesopelagic (800 m, 1000 m) depth zones. Network topologies revealed depth-specific trends of network complexity determined by the total number of nodes (amplicon sequence variants or ASVs), edges (interactions or links), and link density (edge-to-node ratio) (Landi et al., 2018). We explored the influence of complexity across depth zones on network stability, determined by sequentially removing ASVs from the networks (node connectivity and robustness) and edges (edge connectivity) (Iyer et al., 2013; Tipton et al., 2018). These analyses provide a survey of water column biodiversity and reveal insights on the role of keystone taxa and their potential interactions in food webs and ecosystem resilience.

## 3. MATERIALS & METHODS

### 3.1. Sample Collection

Bulk water samples were collected in Northwest Atlantic Slope Water off the coast of New Jersey, eastern United States in March 2020 on the *R/V* Armstrong (AR43) at six discrete depths (10 m, 100 m, 300 m, 500 m, 800 m, and 1000 m) at three sampling stations (Figure 1, Supplementary Table S1). Sampling spanned surface depths to the base of the mesopelagic and included the deep scattering layer (∼500 m) as determined by a broadband Simrad EK80 split beam echosounder system on the *R/V* Armstrong (Supplementary Figure S1, Supplementary Methods). Water was collected with a 24-Niskin CTD rosette across three casts at sampling sites with similar water column characteristics (Figure 1). Two casts (001 and 007) were collected at midnight while cast 002 was collected at mid-day. At each depth, four replicates were collected for a total of 72 samples. For each sample, ∼5 L of water was pumped through sterile 0.2 µm PES Sterivex (MilliporeSigma, Burlington, MA, USA) filters, sealed with Luer lock caps, and stored at −80°C. Sample collection carboys and filtration tubing were bleach-sterilized and rinsed with Milli-Q water prior to filtration with a Masterflex L/S peristaltic pump system following established protocols (Govindarajan et al., 2021, 2022).

**Figure 1.**
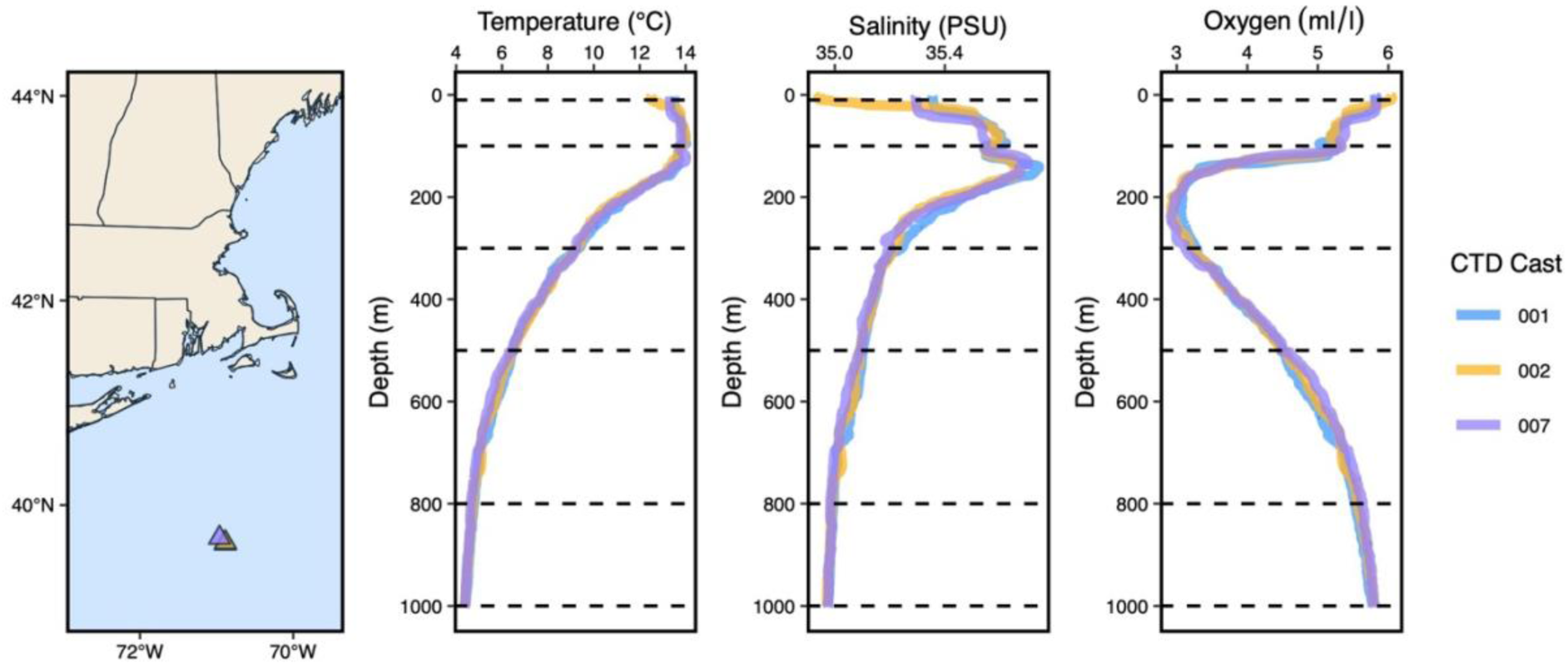
Sampling location and CTD profiles for eDNA collection. Temperature, salinity, and oxygen profiles for each CTD cast (001, 002, and 007) where eDNA samples were collected are shown. Dashed black lines indicate the depths where samples were collected (10 m, 100 m, 300 m, 500 m, 800 m, and 1000 m). Coordinates for sampling sites are provided in Supplementary Table S1.

The AR43 expedition coincided with high densities of the salp *Salpa aspera*, gelatinous zooplankton that are known to periodically “bloom” in the Northwest Atlantic Slope Water (Wiebe et al., 1979; Madin et al., 2006). From analysis of in situ high speed (15 Hz) image data, we estimated densities of 5-10 individuals m^−3^ in surface waters at night (Supplementary Figure S2, Supplementary Methods), which are in the typical range for bloom densities (e.g., 3.5 individuals m^−3^ and higher) (Groeneveld et al., 2020).

### 3.2. DNA Extractions and PCR

DNA was extracted with a Qiagen DNEasy Blood & Tissue extraction kit (Qiagen, Germantown, MD, USA) following manufacturer’s protocols with modifications for Sterivex filters (Govindarajan et al., 2021). Extracted DNA was stored at −20°C prior to metabarcoding. All samples were metabarcoded for portions of the 18S and 12S rRNA genes (see below), except for one sample collected on CTD cast 007 at 1000 m which did not have enough DNA template for 12S rRNA metabarcoding. All lab surfaces and equipment were cleaned with a 10% sodium hypochlorite solution (household bleach) and extractions were performed in a dedicated laboratory space away from sources of concentrated DNA such as post-PCR or tissue samples.

The 18S V9 hypervariable region was amplified using the 1380F/1510R primer set with CS1/CS2 Fluidigm adapters and following a two-step PCR protocol (Amaral-Zettler et al., 2009; Govindarajan et al., 2021). The initial PCR cycling program began with a denaturation step at 95°C for 3 min, 25 cycles at 95°C for 30 sec, 55°C for 30 sec, and 72°C for 30 sec, followed by an elongation step at 72°C for 5 min. PCR reactions (25 μl) consisted of 2.5 μl of DNA template, 0.5 μl of each primer, 12.5 μl of 2x KAPA HiFi HotStart ReadyMix (Kapa Biosystems; Wilmington, MA, USA), and 9 μl of molecular-grade water. Duplicate PCR reactions were run for each sample, pooled, and sent to the Genome Research Core at the University of Illinois at Chicago (UIC) to complete a second PCR amplification to add unique tags using MyTaq HS 2X Master Mix and the Access Array Barcode Library for Illumina (Fluidigm, South San Francisco, CA, USA). The PCR cycling program was as follows: 95°C for 5 min followed by 18 cycles of 95°C for 30 sec, 60°C for 30 sec, 72°C for 30 sec, and 72°C for 7 min. Amplicon libraries were purified using AMPure XP beads (Beckman Coulter, Indianapolis, IN, USA). Amplicon sequencing with a targeted sequencing depth of 100000 reads per sample was done on an Illumina MiSeq (2×250 bp) (Govindarajan et al., 2022).

The hypervariable 12S region of the mitochondrial rRNA gene was amplified using MiFish primers (Miya et al., 2015) appended with Illumina adapter overhang nucleotide sequences and a modified two-step PCR protocol with a touch-down PCR program for the first PCR (Pitz et al., 2020). The initial PCR cycling program began with a denaturation step at 95°C for 15 min; 13 cycles of 94°C for 30 sec, 69.5°C for 30 sec (decreasing 1.5°C with each cycle), and 72°C for 90 sec; 25 cycles of 94°C for 30 sec, 50°C for 30 sec, and 72°C for 45 sec; followed by an elongation step at 72 °C for 10 min. PCR reactions (25 μl) consisted of 2.5 μl of DNA template, 1 μl of each primer, 12.5 μl of 2x KAPA HiFi HotStart ReadyMix, and 8 μl of molecular-grade water. Duplicate PCR products were pooled and sent to the Center for Genome Innovation at the University of Connecticut for size selection, a second round of PCR, and sequencing. Amplicons from the pooled PCR products were visualized with the Agilent 4200 TapeStation electrophoresis system using the High Sensitivity DNA D1000 assay (Agilent Technologies, Santa Clara, CA). Size - selection for targeted amplicons (215bp-335bp) was conducted using the Sage Science PippinPrep HT gel cassette (2% Agarose, PippinHT). Illumina barcodes were added to each amplicon using a second PCR amplification. The PCR cycling program was as follows: 95°C for 3 min followed by 12 cycles of 95°C for 15 sec, 60°C for 30 sec, and 72°C for 90 sec. Amplicon libraries were purified using AMPure XP beads and assessed for quality and adapter dimer content on the Agilent 4200 TapeStation using a D1000 assay. Amplicon sequencing with a targeted sequencing depth of 100000 reads per sample was done on an Illumina MiSeq (2×250bp) with 20–30% PhiX added (Illumina, San Diego, CA).

For both 18S and 12S primer sets, PCR runs included field controls of filtered Milli-Q water (1 L for each CTD cast) as well as PCR negative (without DNA template) and positive controls. The PCR positive control was included to determine the quality of sequencing results and consisted of previously extracted 1ng/μl sample of *Fundulus heteroclitus*, an estuarine fish that would not be found in the environmental samples.

### 3.3 Multi-marker metabarcoding bioinformatics

Demultiplexed sequencing reads were imported into and processed in QIIME 2 with default settings (v2022.11.1) (Bolyen et al., 2019). Adapters and primers were trimmed from the reads with Cutadapt (Martin, 2011). Reads were quality-filtered, trimmed to the primer region 120 bp (18S) and 150 bp (12S), denoised, and merged into ASVs with DADA2 (Callahan et al., 2016). ASVs represent exact biological sequences that can be distinguished by as little as one nucleotide (Callahan et al., 2017). For both gene markers, taxonomy was assigned by a naïve Bayes classifier at an 80% confidence threshold (Pedregosa et al., 2011); the classifier was trained on reference databases curated and evaluated with the REference Sequence annotation and CuRatIon Pipeline (RESCRIPt) QIIME 2 plugin (Robeson et al., 2021).

The 18S sequencing reads were annotated with two different databases: one for protists and one for metazoans. The protist (18S) classifier was trained with the Protistan Ribosomal Reference (PR2) database (v5.0.1) (Guillou et al., 2013) that was updated with 9 taxonomic levels including the replacement of kingdom with domain and the addition of a new subdivision level. The metazoan (18S) classifier was trained with the SILVA SSURef_NR99 (v138.1) database (Pruesse et al., 2007; Quast et al., 2013). A new 12S classifier was created by combining a custom reference database of 12S sequences for 80 species of mesopelagic fish from the Northwest Atlantic (Govindarajan et al., 2023b) with the MetaZooGene database (v2023-m07-15) in Mode-A (most inclusive option) for vertebrates (O’Brien et al., 2024). For 18S sequences, taxonomy assignments were first assigned with PR2 and protist sequences were selected as the subset where phylum was not classified as “Metazoa”. This subset will be referred to as the protist dataset. Sequences where phylum was classified as “Metazoa” were subsequently classified with the SILVA-based classifier. Metazoan 18S reads were classified predominantly as invertebrate taxa and will be referred to as the invertebrate dataset (Govindarajan et al., 2021). Likewise, metazoan 12S reads were classified as vertebrates (mainly fishes) and will be referred to as the vertebrate dataset. For downstream bioinformatic and statistical analyses, the 18S reads were processed separately as two individual datasets representing protist and invertebrate reads, while 12S reads represented the standalone vertebrate dataset.

After taxonomic classification, 142 of 399 invertebrate ASVs remained unassigned at the phylum level or were unassigned sequences from the phylum Arthropoda, which represented 18.4% of all sequences in the invertebrate dataset. Of those 142 ASVs, one ASV represented ∼13% of invertebrate reads and comprised of ∼48% of reads at 100 m (Supplementary Figure S3A). These unassigned sequences were manually classified with Geneious Prime 2025.0.3 with a BLAST search and NCBI’s new BLAST core nucleotide database (core_nt). The top match was selected from the assigned grade score at a 95% cut-off. This approach annotated 67 of the 142 unassigned invertebrate ASVs, which reduced the total number of unassigned reads to 0.35%. Most unassigned reads were classified to Order Calanoida (calanoid copepods). Similarly, 345 of 677 vertebrate ASVs remained unassigned with > 80% of ASVs classified as Unassigned Actinopterygii (ray-finned fishes). Altogether, unassigned ASVs comprised 12.9% of reads. Following the BLAST process described for invertebrates, 117 vertebrate ASVs were successfully classified, reducing the unassigned read percentage to ∼1% (Supplementary Figure S3B).

Taxonomy, ASV count, and metadata files were imported into R (v4.3.2) with qiime2R (v0.99.6) (https://github.com/jbisanz/qiime2R) and merged with phyloseq (v1.46.0) (McMurdie and Holmes, 2013) to generate phyloseq objects for protists, invertebrates, and vertebrates. Potential contaminants were identified and removed with the R package decontam and a conservative threshold of 0.5, which identifies all sequences more prevalent in negative controls than samples as contaminants (Davis et al., 2018). In total, decontam identified and removed 44 protist, 19 invertebrate, and 17 vertebrate ASVs. Additional decontamination of the datasets was conducted manually and detailed in Supplementary Methods. Sequencing thresholds were selected to remove low read-depth outliers (Schloss, 2024a) such that samples with less than 15000 reads (protists), 6000 reads (invertebrates), and 3000 reads (vertebrates) were removed, resulting in 69 samples (protists), 70 samples (invertebrates), and 61 samples (vertebrates) out of 72 total samples (Supplementary Figure S4).

### 3.4 Community Ecology Analyses

Alpha diversity indices (richness, Shannon Index, evenness) were calculated from a rarefaction approach with iterative subsampling (1000 iterations) and a subsampling threshold (Schloss, 2024a) determined by the minimum read count of 27865 (protists), 6076 (invertebrates), and 3183 (vertebrates) in the sample set (Schloss, 2024b). Diversity indices reflect the mean index value for each depth after rarefaction and statistical significance was computed by one-way ANOVA with Tukey’s HSD post-hoc analysis (*p* value < 0.05).

Depth-specific community composition was assessed with principal component analyses conducted with the rda function in vegan v2.6.4 (Oksanen et al., 2022) on centered log-ratio (CLR) transformed ASV counts (Aitchson, 1986; Quinn et al., 2018; Coenen et al., 2020). Zero-value data were replaced with the “const” method which replaces zeros with 65% of the next lowest value (Bastiaanssen et al., 2022). Statistical significance was assessed with permutational multivariate analysis of variance (PERMANOVA) with the adonis2 function in vegan on Euclidean distance matrices calculated from CLR-transformed data (i.e., Aitchison distance). Pairwise comparisons were conducted with the pairwise.adonis2 function from the pairwiseAdonis package v0.4 (Arbizu, 2020).

ASVs were grouped into relevant taxonomic groups for taxa bar plots to show the proportion of ASV composition and relative abundance. UpSet plots were used to visualize ASV overlap across depths, with a matrix format that shows which ASVs are shared or unique to specific depths. Taxa with low relative abundance (< 1%) were grouped to streamline visualizations with additional details provided in Supplementary Materials (Supplementary Tables S2-S4). ASV accumulation curves were generated with the specaccum function in vegan v2.6.8 with the “random” method, which randomizes sample order and averages ASV richness across 999 permutations to show how ASV richness increases with sampling effort.

### 3.5 Network Analyses

Network analysis from Sparse InversE Covariance Estimation for Ecological Association inference (SPIEC-EASI) was applied to protist, invertebrate, and vertebrate compositional datasets to assess ASV-ASV co-occurrence patterns with depth across these datasets (Kurtz et al., 2015; Tipton et al., 2018). SPIEC-EASI identifies statistically significant relationships while accounting for issues inherent to compositional metabarcoding datasets (Kurtz et al., 2015). ASV count tables are CLR-transformed and a graphical model inference framework is used to identify direct associations between ASVs while minimizing indirect correlations and reducing the detection of spurious relationships.

Samples were grouped into three depth zones: epipelagic (10 m and 100 m), upper mesopelagic (300 m and 500 m), and lower mesopelagic (800 m and 1000 m). Out of a possible total of 24 samples in each depth zone, only samples that contained protist, invertebrate, and vertebrate ASVs were retained for network analysis, resulting in a total of 15 epipelagic samples, 21 upper mesopelagic samples, and 22 lower mesopelagic samples. ASVs were filtered on abundance (> 50 reads) and prevalence (present in at least 10% of samples in each depth zone) to minimize dense networks and focus the analyses on prevalent and abundant ASVs.

Undirected networks were generated for each depth zone for metazoan-only (invertebrate and vertebrate) ASVs, protist-only ASVs, and all ASVs (All Taxa) for a total of nine networks. The SpiecEasi v1.1.1 package was used to construct these networks on WHOI’s high-performance computing cluster in R v4.4.3, applying the Meinshausen–Buhlmann’s neighborhood selection method with an optimal sparsity threshold (lambda min ratio) of 1e-3, and 100 subsampling replicates (rep.num = 100) (Kurtz et al., 2015).

In each network, edges (links) were inferred using SPIEC-EASI and represent signed conditional associations between ASVs (nodes) after accounting for indirect associations mediated by other ASVs. Positive edges indicate ASVs that exhibit similar relative abundance patterns across samples (co-vary in the same direction), whereas negative edges indicate ASVs that co-vary in opposite directions

Networks were then exported into RStudio to assess network topology and robustness with igraph v2.0.2 in R (Csárdi and Nepusz, 2006). For each network, we evaluated node and edge connectivity, which represents the minimum threshold for node or edge removal that fragments a connected network. We also evaluated attack robustness, which measures the size of the network (remaining subnetwork) as nodes are sequentially removed based on betweenness, degree, and at random (Iyer et al., 2013; Tipton et al., 2018).

Although ecological networks can be built from species-level or other taxonomic groups, we built networks from ASVs, the lowest genetic unit, to render an appropriate comparison across the three datasets generated from two markers of varying taxonomic resolution. For example, while 18S V9 is highly conserved and is useful for broad amplification of eukaryotic biodiversity, it has less resolving power than other markers like 12S MiFish, which can resolve to species level for a wide variety of fishes (Creer et al., 2016; Zhang et al., 2018; Miya et al., 2020; Govindarajan et al., 2021; Questel et al., 2021). To show this, we evaluated the three naïve Bayes classifiers referenced in this study with the evaluate-fit-classifier function in the QIIME 2 RESCRIPt plugin (Robeson et al., 2021) to test the classifiers’ performance on the same sequences it was trained with. The F-measure score, which reflects the balance between precision and recall, for the 18S V9 classifiers drops below ∼0.95 after Order while the 12S classifier has a high F-measure up to species-level classification (Supplementary Figure S5). After networks were generated, ASVs were grouped into relevant taxonomic groups of ecological and functional significance for downstream network analyses.

### 3.6 Visualizations

Unless otherwise specified, figures were generated with ggplot2 v3.5.1 (Wickham, 2016). UpSet plots were generated with ComplexUpset v1.3.3. Networks were visualized with ggnetwork v0.5.13 (Tyner et al., 2017). When necessary, visualizations were compiled with minimal editing for aesthetics with the creative software, Affinity Designer 2 (Serif, Europe).

## 4. RESULTS

### 4.1 Patterns of diversity from 18S and 12S metabarcoding

Metabarcoding of the 18S V9 rRNA gene generated ASVs for protists and invertebrates while the 12S MiFish metabarcoding generated ASVs for vertebrates (predominately fishes). The 18S dataset ranged from 70-90% protist sequences with the proportion of invertebrate sequences decreasing with depth (Supplementary Figure S6). Protist and invertebrate sequences were parsed from the 18S dataset and assessed separately. Following decontamination and rarefaction of metabarcoding reads, a total of 8112 ASVs were recovered across both markers with 7046 (86.9%) for protists, 399 (4.9%) for invertebrates, and 667 (8.2%) for vertebrates. The numbers of observed ASVs (or ASV richness) for protists and invertebrates were generally higher in the mesopelagic than the epipelagic with relatively constant richness across mesopelagic depths (Figure 2). For vertebrates, higher richness was observed at 10 m and 500 m than at other depths. For invertebrates, Shannon diversity indices were significantly different between surface (10 m) and all other depths (*p* < 0.01) while Pielou’s evenness indices varied significantly at depth, especially between 100 m and select mesopelagic depths (*p* < 0.05). For vertebrates, average Shannon and Pielou’s indices followed similar patterns as observed for ASV richness, although no significant differences were observed between depths (Supplementary Tables S5-S7). For protists however, Shannon and Pielou indices exhibited patterns that contrasted with those for richness. The highest Shannon diversity and Pielou evenness indices occurred at 100 m and then decreased with depth down to 500 m before increasing again to 1000 m.

**Figure 2.**
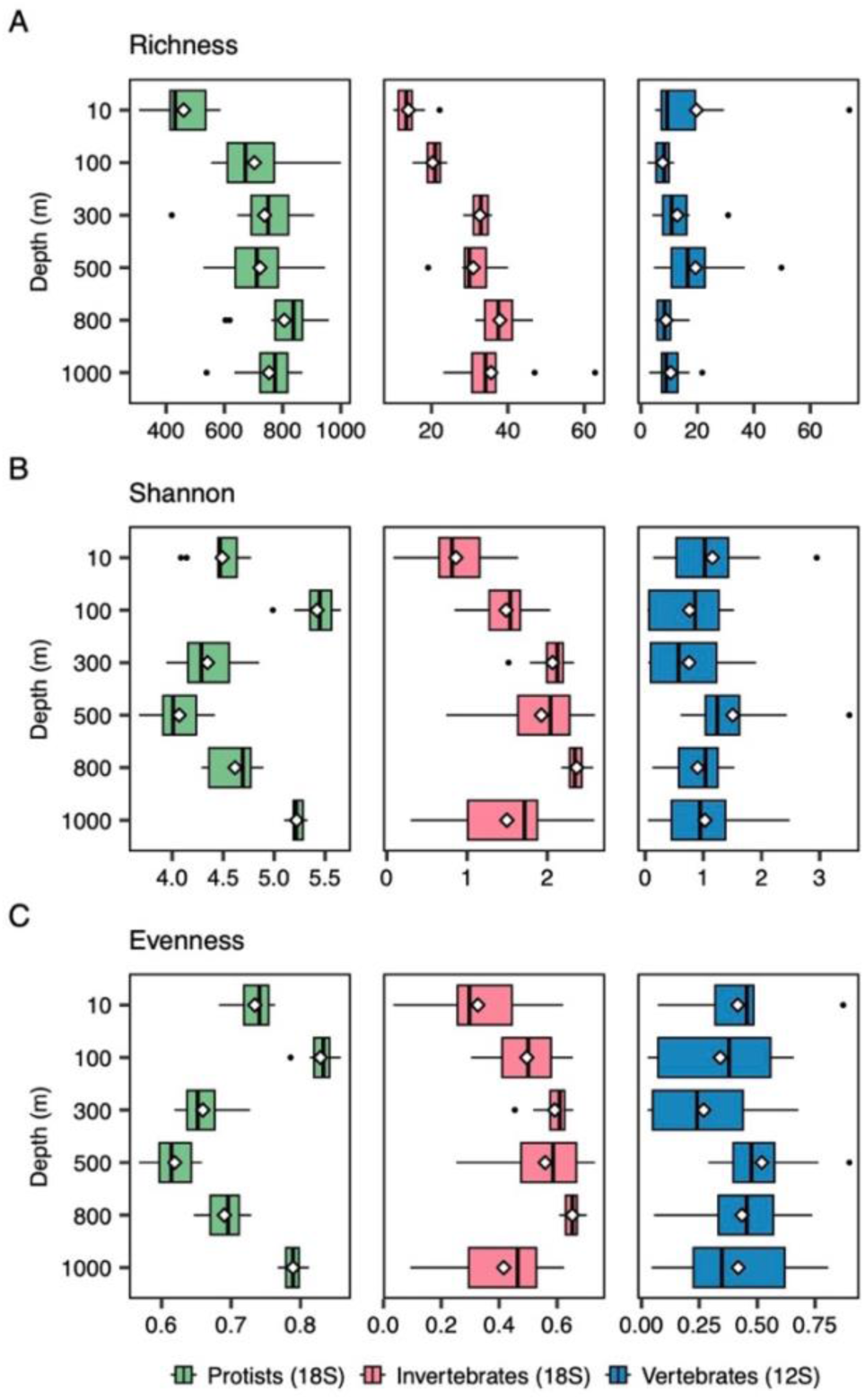
Patterns of alpha diversity with depth for A) Richness (observed ASVs), B) Shannon Diversity Index, and C) Pielou’s Evenness Index across all samples for protists (n=69), invertebrates (n=70), and vertebrates (n=61). Boxplots show the variation in diversity indices at each depth between samples divided into quartiles with outlier values indicated by black points. A white diamond indicates the mean value at each depth.

Protistan diversity in epipelagic waters was comprised of mostly alveolates (dinoflagellates, Syndiniales, and ciliates) and stramenopiles (e.g., diatoms) (Supplementary Figure S7A). Alveolate ASVs increased in mesopelagic waters, as did rhizarian (e.g., radiolarians) and excavate ASVs, with these three groups representing a significant portion of protistan diversity (Supplementary Figure S7D). Invertebrate diversity was driven mainly by taxonomic groups with relative read abundance < 1% (rare reads), including Trachymedusae and Narcomedusae (Cnidaria) (Supplementary Figure S7B, Supplementary Table S3). Orders Calanoida (calanoid copepods) and Siphonophorae (Cnidaria; siphonophores), as well as unassigned metazoan ASVs, also represented a significant portion of diversity (Supplementary Figure S7E). Fish were the main category of vertebrate ASVs detected (Supplementary Figure S7C), with a significant portion attributed to deep sea fish taxa including Melamphidae (bigscales), Myctophidae (lanternfish), and Gonostomatidae (bristlemouths). Similar to the invertebrates, rare taxonomic groups (low read count) contributed significantly to vertebrate diversity (Supplementary Figure S7F). Five mammalian ASVs were also detected, comprising ∼0.038% of the total dataset. These included ASVs assigned to whales and dolphins known to occur at the study site, including two ASVs from the genus *Balaenoptera* (whales) and three ASVs for dolphins in the genera *Delphinus*, *Stenella*, and *Grampus* (Supplementary Table S4).

### 4.2 Depth structuring of the eukaryotic community

Principal component analysis (PCA) and PERMANOVA of rarefied, center-log ratio (CLR) transformed protist and invertebrate reads from all samples revealed significant depth clustering (*p* < 0.001) (Figure 3). Moreover, pairwise comparison showed that communities at each depth were significantly different from other depths (*p* < 0.001). For vertebrate communities, significant differences were observed between surface (10 m) and mesopelagic depths (*p* < 0.01). Differences were also found between 500 m and all other depths (*p* < 0.05). When all depths are considered, clustering by CTD cast, which reflected the time of sampling, was not statistically significant (*p* > 0.05). Given the limited number of CTD casts (two night and one day) and the lack of significant diel patterns in this dataset, samples across all three casts were combined at each depth for subsequent analyses (Supplementary Figures S8-S9, Supplementary Discussion).

**Figure 3.**
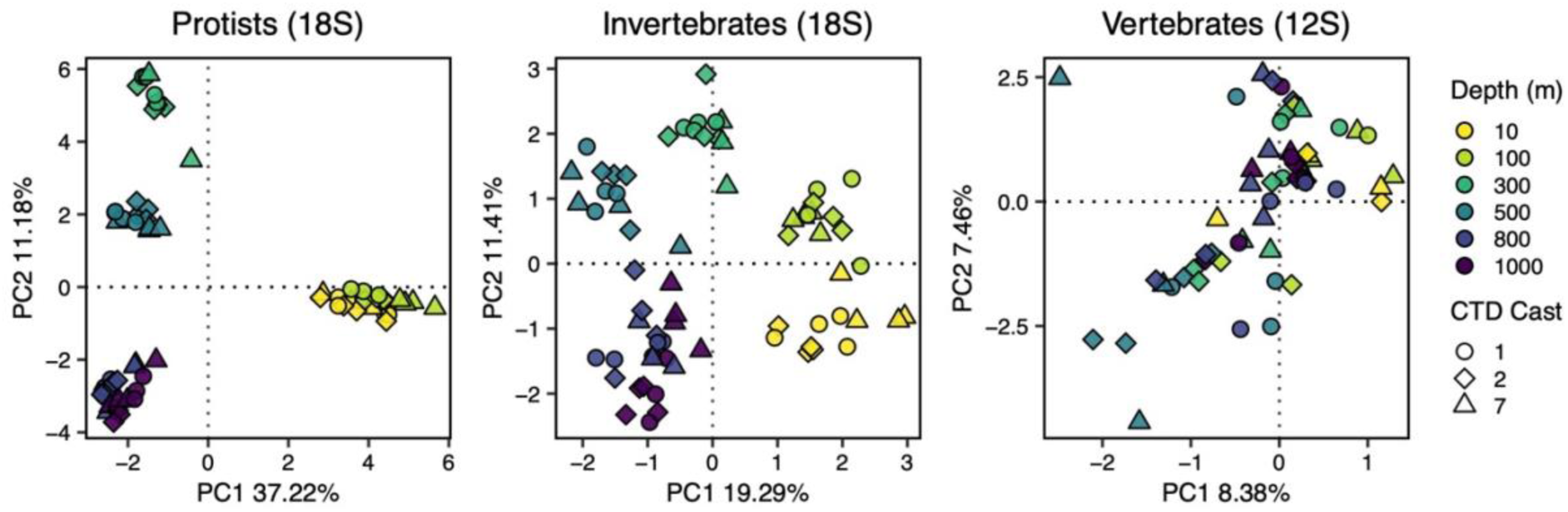
Principal component analysis (PCA) of center-log ratio-transformed reads for protists (n=69), invertebrates (n=70), and vertebrates (n=60). Each point represents one sample. The shape and color of each point indicates the CTD cast and the sampling depth, respectively, of each sample. CTD cast here can also reflect diel sampling with casts 1 and 7 sampled at midnight and cast 2 sampled during mid-day. One outlier (10 m, PC1 = ∼10) from the vertebrate dataset was removed for visualization purposes (see Supplementary Figure S10).

Sequenced reads were aggregated by depth to generate relative abundance profiles that reflect the overall water column community structure (Figure 4). For protists, epipelagic (< 200 m) community structure was dominated by photosynthetic taxa including dinoflagellates, diatoms and other stramenopiles, and chlorophytes. The relative abundance of the parasitic taxon Syndiniales, as well as radiolarians, increased with depth and were the dominant taxa in the mesopelagic (> 200 m). Excavates were also relatively more abundant in deeper waters. For invertebrates, calanoid copepods (Arthropoda) exhibited higher relative abundance in both the epipelagic and lower mesopelagic (1000 m), while siphonophores (Cnidaria) were also relatively abundant, especially in the upper mesopelagic. At surface depths, salps were relatively abundant compared to other invertebrate taxa.

**Figure 4.**
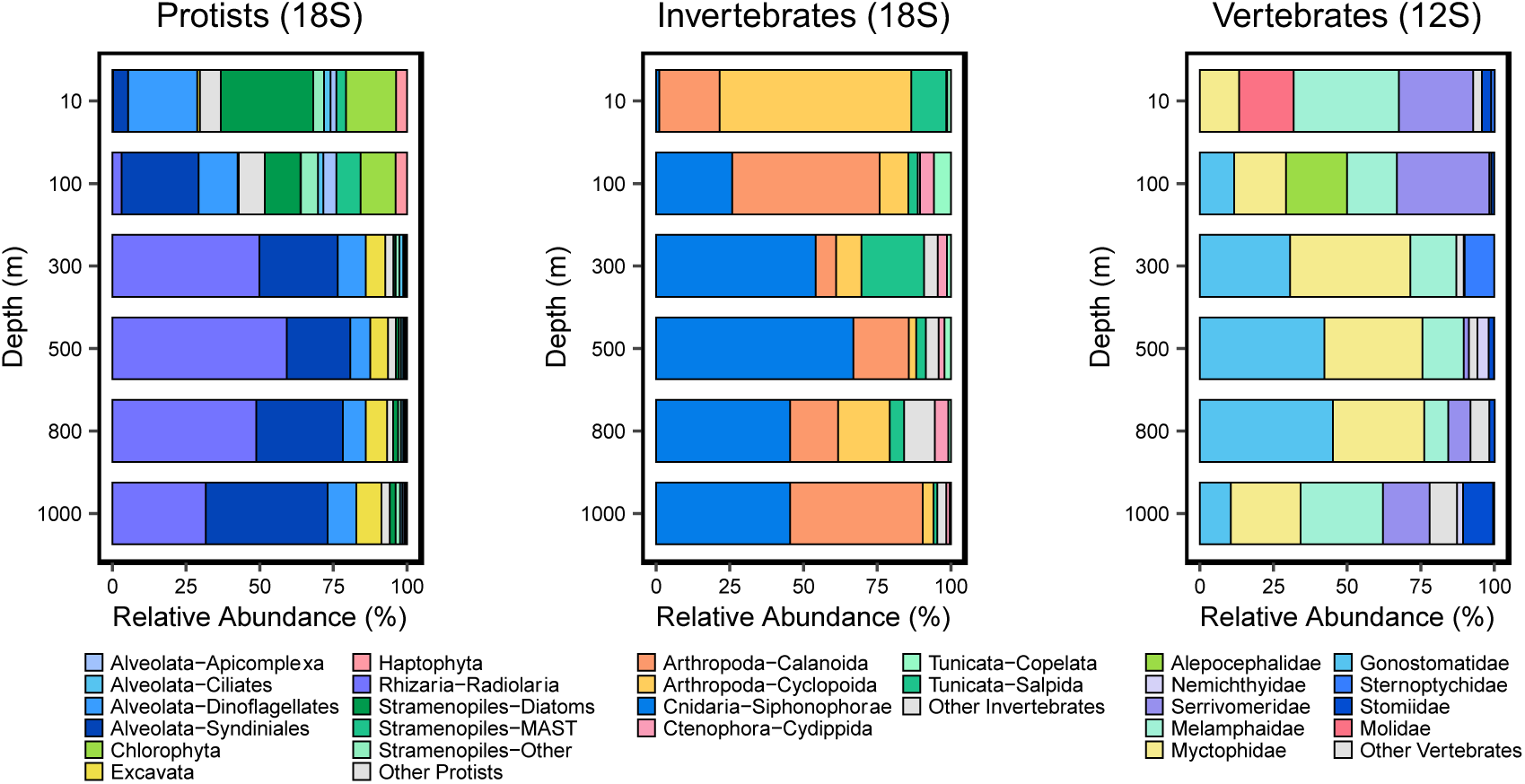
Depth profiles of relative read abundance (%) for protists, invertebrates, and vertebrates calculated from samples aggregated across three CTD casts. Bar plots show taxonomic groups in the order of decreasing relative abundance. In the legend, taxonomic groups within protists (Supergroup and/or Division or Subdivision) and invertebrates (Phylum and Order) are colored and listed in alphabetical order. Vertebrate families were grouped at the Order level in alphabetical order: Alepocephalidae (Alepocephaliformes); Nemichthyidae, Serrivomeridae (Anguilliformes); Melamphidae (Beryciformes); Myctophidae (Myctophiformes); Gonostomatidae, Sternoptychidae, Stomiidae (Stomiiformes); and Molidae (Tetraodontiformes). For each group, only taxa with overall relative abundance > 1% are displayed, with all other taxa combined in “Other”. For a full list of “Other” taxa, see Supplementary Tables S2-S4.

Key taxonomic groups were comprised of ASVs that were only detected at one specific depth (unique ASVs), detected at two or more depths (shared ASVs), or detected at all depths (cosmopolitan ASVs) (Supplementary Figure S11). Unique, shared, and cosmopolitan protist ASVs included a mix of both photosynthetic and non-photosynthetic taxa across all depths with deeper waters harboring a higher number of unique ASVs than near the surface (10 m). Unique invertebrate ASVs were greater at mesopelagic depths and dominated by taxa with low relative abundance (< 1%), while unique vertebrate ASVs were highest at 10 m and 500 m. Across the three organismal groups, unique ASVs were proportionally higher than shared and cosmopolitan ASVs, especially for invertebrates (68.4%) and vertebrates (89.2%) compared to protists (56.6%).

For invertebrates, shared and cosmopolitan ASVs included representatives from Orders Calanoida (Arthropoda), Siphonophorae (Cnidaria), Cydippida (Ctenophora), as well as Copelata and Salpida (Tunicata) (Supplementary Figure S12). For vertebrates (Supplementary Figure S13), shared and cosmopolitan ASVs were observed from mesopelagic fish species including *Benthosema glaciale* (Myctophidae), *Cyclothone microdon*, (Gonostomatidae), *Scopeloberyx opisthopterus* (Melamphaidae), and *Serrivomer beanii* (Serrivomeridae).

### 4.3 Depth-specific eukaryotic networks

We generated single-taxon and multi-trophic networks of protists, invertebrates, and vertebrates for the epipelagic (10 m and 100 m), upper mesopelagic (300 m and 500 m), and lower mesopelagic (800 m and 1000 m). Associations between “Metazoa-only” (invertebrates and vertebrates), “Protists-only”, and All Taxa (all groups) for each depth zone were inferred with SPIEC-EASI to assess the contribution of broad eukaryotic groups to network connectivity and stability.

Metazoa-only networks were small and fragmented, consisting of multiple components including singleton and doubleton associations that resulted in lower complexity networks compared to Protist-only and All Taxa networks (Figure 5, Supplementary Table S8). In contrast, Protist-only networks and All Taxa networks were fully connected such that a link existed between any two nodes in the network. Across all networks, the epipelagic had lower complexity (fewer nodes and edges) and decreased edge density (edge-to-node ratio) compared to the mesopelagic depths.

**Figure 5.**
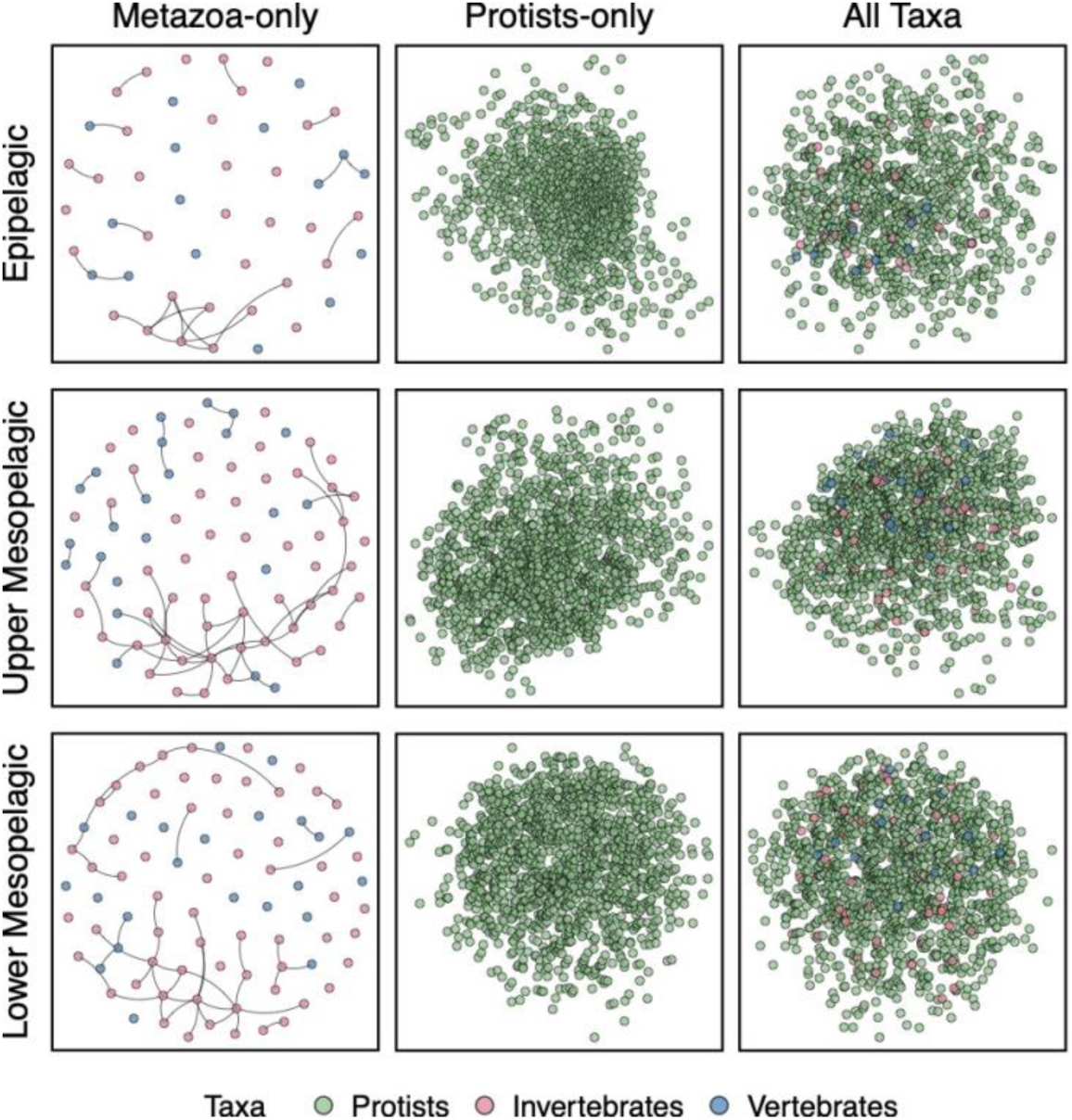
Depth-specific networks for Metazoa-only (invertebrate and vertebrates), Protists-only, and All Taxa (all ASVs). All nodes are shown as points with color indicating the broad taxonomic classification of protists, invertebrates, and vertebrates. Edges are shown for the Metazoa-only network to show the fragmented nature of the network with singleton nodes. The Protists-only and All Taxa networks are connected networks and edges are not shown. All networks include predicted positive and negative interactions. A larger version of this figure is available as Supplementary Figure S15.

For each network, we assessed resilience and stability to the loss of ASVs. Node and edge connectivity increased with depth for All Taxa networks (Supplementary Table S8). The lower mesopelagic had the highest threshold for node and edge connectivity for both All Taxa and Protist networks. For the already fragmented Metazoa-only networks, connectivity was zero across all depth zones. We also evaluated attack robustness by which measuring the size of the remaining subnetwork as nodes are sequentially removed based on betweenness (putative bottlenecks or keystone taxa), degree (highly connected ASVs or hubs), and at random (Iyer et al., 2013; Tipton et al., 2018). Metazoa-only networks were the least robust, especially in the mesopelagic. Protists-only and All Taxa networks had comparable robustness with higher robustness in mesopelagic networks (Supplementary Figure S14).

### 4.4 Ecological interactions from the surface to the mesopelagic

Positive and negative interactions were separated from the combined networks to explore putative ecological interactions, especially trophic dynamics across depth. While these networks do not represent direct ecological interactions, associations between ASVs can be used to hypothesize potential ecological relationships among taxa (Landi et al., 2018). Epipelagic and upper mesopelagic networks averaged 58 and 57% positive interactions, respectively, while the lower mesopelagic network was 53% positive (Supplementary Table S9). On average, the edge density of positive networks was 8.1 compared to 6.5 for negative networks, suggesting that positive networks are more complex. Nodes were compartmentalized into modules that delineated subsets of ASVs that interact more frequently or have stronger interactions than with other ASVs (modularity). Average modularity across the three depth zones was higher for positive networks compared to negative networks (Supplementary Table S10). For both positive and negative networks, modularity decreased with depth. Node and edge connectivity for negative networks averaged 1.33 compared to an average of 2.67 for positive networks. Negative networks were also less robust than positive networks, especially in the epipelagic networks (Supplementary Figure S16).

Nodes with the highest edge weights (90^th^ percentile), indicating the strongest interactions, were isolated from the positive and negative networks to explore key ASV associations in each network (Supplementary Figure S17). In both positive and negative subnetworks, the top ranking ASVs based on node degree (number of connections) and betweenness centrality were predominantly protistan taxa, especially Syndiniales (Alveolata) (Supplementary Figures S18-S19). Top ranking invertebrate ASVs included copepods (both calanoid and cyclopoid), siphonophores (cnidarians), and ctenophores while vertebrate ASVs included mesopelagic fish. Notably, *Grampus griseus* (Risso’s dolphin) was a top-ranking ASV in the positive network, while salps (tunicates) and *Coryphaena hippurus* (common dolphinfish) were high-ranking ASVs in the negative network.

Trophic interactions were assessed by assuming a simple food web from the negative network where protists are at the bottom, invertebrates are intermediate, and vertebrates are top predators. In general, the total number of pair-wise associations (protist-invertebrate, protist-vertebrate, and invertebrate-vertebrate) were highest in the mesopelagic, with 1193 and 1088 associations in the upper and lower mesopelagic, respectively, compared to 582 associations in the epipelagic (Figure 6). We narrowed focus to the strongest interactions at each depth zone by selecting ASV-ASV associations with the top 10% of edge weights. Many of these strongest relationships featured protist associations with both invertebrates and vertebrates and highlighted known interactions such as zooplankton grazing of phytoplankton, predator-prey dynamics between zooplankton and mesopelagic fish, as well as putative parasitic interactions between alveolates and various metazoans (Figure 6).

**Figure 6.**
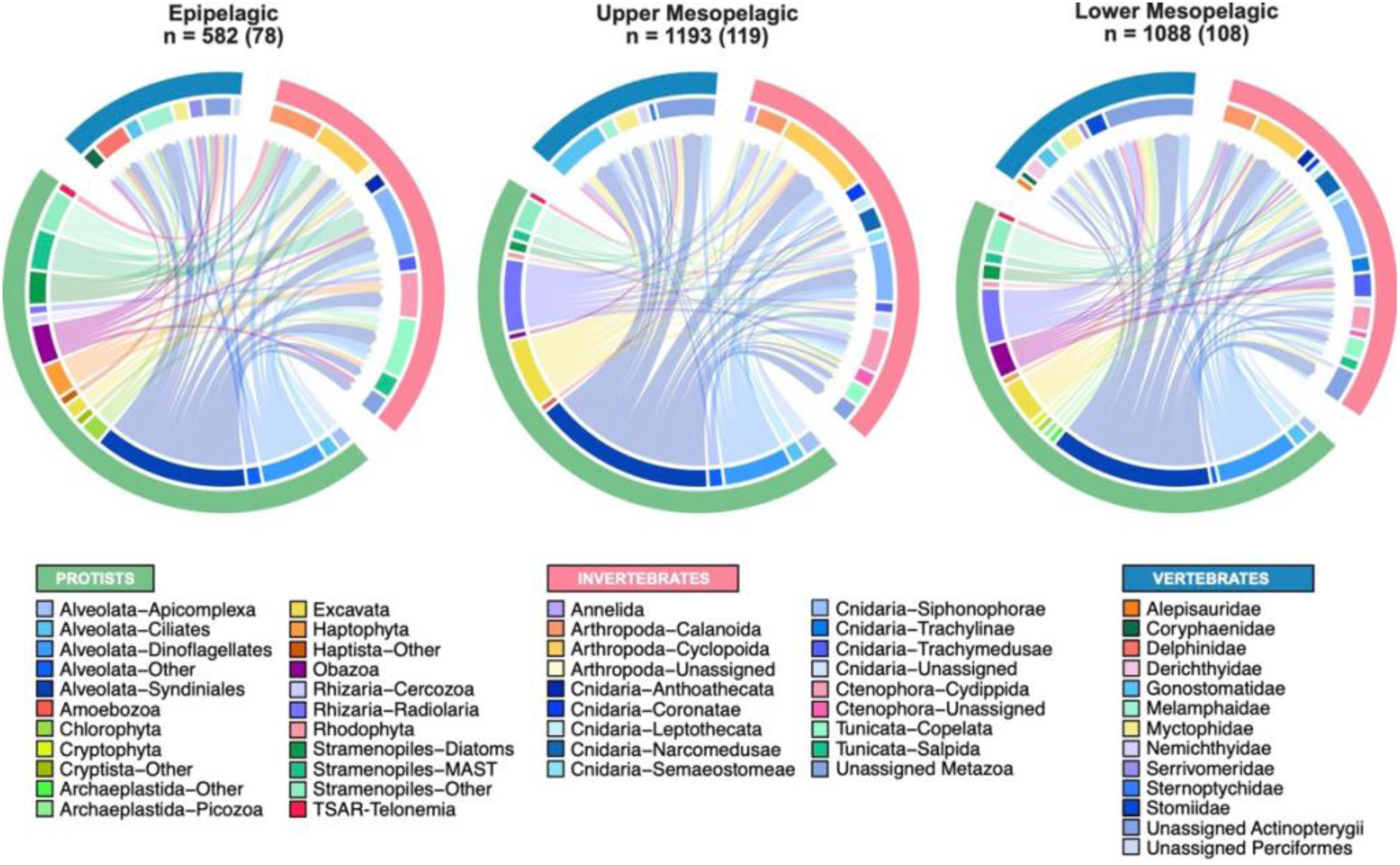
Chord diagrams indicate the number of strong pairwise interactions in the negative network and the taxonomic groups involved at each depth zone. Interactions were selected by removing all ASV-ASV linkages below the 90^th^ percentile threshold for edge weight (so that only the strongest interactions remained). Only protist-invertebrate, protist-vertebrate, and invertebrate-vertebrate interactions were considered. The total number of pairwise associations for each negative network is listed (n) with the number of strong associations in parentheses. Colors along the circumference (track) of the diagram indicate taxonomic groupings. The outer track shows protists in green, invertebrates in pink, and vertebrates in blue. The inner track further divides these groupings into taxonomic groups. Each link between two ASVs (or interaction) is depicted as a directional arrow that starts from the ASV in a lower trophic level and points towards the ASV at the higher trophic level. The arrows are also colored based on the lower trophic level ASV. Trophic levels are simplified and ordered as protist > invertebrate > vertebrate. The size of each inner track segment indicates the relative number of ASVs under each taxonomic group within a chord diagram. A larger version of this figure is available as Supplementary Figures S20-S22.

## 5. DISCUSSION

### 5.1 Protists and metazoans exhibit contrasting diversity patterns

Based on current knowledge of the deep sea, we lack consensus on broad diversity trends due to prevailing gaps across spatial, temporal, and taxonomic scales (Haddock and Choy, 2024; Yang et al., 2024). Traditional midwater assessments often rely on net-based trawls, which sample few depths and regions and can underrepresent taxa that avoid capture or are too fragile (Kaartvedt et al., 2012; Gold et al., 2021; Suarez-Bregua et al., 2022). As such, deep sea studies that span the eukaryotic community from microeukaryotes to metazoans remain scarce. Since we applied multi-marker metabarcoding to eDNA, this study has generated new insights into eukaryotic biodiversity and ecology from the surface down to the base of the mesopelagic.

A key finding of our study is that alpha diversity increased in the mesopelagic compared to the epipelagic. We found this for both ASV diversity (richness) and Shannon diversity in the Northwest Atlantic Slope Water, especially for protists and invertebrates. This observation differs from the existing perspective of decreasing diversity in midwater ecosystems compared to epipelagic waters generated from the global Ocean Biodiversity Information System (OBIS) database (Costello and Chaudhary, 2017). However, Costello and Chaudhary did note a diversity maximum at ∼400-500 m, which aligns with our findings, especially for vertebrates (Figure 2). A recent survey of more than two decades of ROV-generated data revealed taxa-specific patterns in the eastern North Pacific such as higher species richness for crustaceans and fishes in the mesopelagic compared to the epipelagic (Haddock and Choy, 2024).

While our findings generally show increasing alpha diversity with depth (Figure 2), we observed low indices for protists between 300 m and 800 m with a minimum at 500 m. We attributed the lower diversity in this depth zone to a small number of ASVs associated with Syndiniales (Dinoflagellata) and Radiolaria (Rhizaria) that are highly abundant relative to other ASVs (Supplementary Figure S23). Through the upper mesopelagic, this trend is generally in line with previous studies that reported a protist diversity maximum at the base of the euphotic zone followed by decreasing diversity in the mesopelagic (Laroche et al., 2020b; Blanco-Bercial et al., 2022). In contrast, other studies have found increasing diversity in the mesopelagic, which we also observed in our dataset at 1000 m (Giner et al., 2020; Ollison et al., 2021).

We also found higher richness in the mesopelagic for invertebrates, which is similar to diversity metrics reported for other areas of the North Pacific (Matthews and Blanco-Bercial, 2023) as well as the southeast Pacific (González et al., 2023), but contrasting with diversity patterns reported off Hawaii (Laroche et al., 2020b). These studies showed that arthropods (especially copepods) and cnidarians are significant contributors to metazoan diversity. In addition, we found vertebrate diversity assessed with the 12S biomarker detected mainly fish. As with previous studies (Canals et al., 2021; Govindarajan et al., 2023b), a significant portion of sequencing reads were associated with deep-sea fish (Supplementary Figure S7).

For invertebrates and vertebrates, Shannon diversity was high across mesopelagic depths with the highest ASV richness at 500 m for vertebrates, which corresponded with the deep scattering layer detected at 38 kHz (Supplementary Figure S1), a region of the midwater with strong acoustic backscatter from mesopelagic fish and invertebrates (Barham, 1966; Aksnes et al., 2017). Taxonomic groups with low relative abundance (< 1%) or “rare” taxa typically contributed to alpha diversity for protists, invertebrates, and vertebrates (Supplementary Figure S7 and S11). While our study was not explicitly focused on the role of these low abundance taxa, these rare taxa may exert considerable influence over ecosystem function and stability (Lyons et al., 2005; Mouillot et al., 2013; Logares et al., 2014; Lynch and Neufeld, 2015; Jousset et al., 2017).

There are various factors that may contribute to the differences reported across metabarcoding studies. There are differences in methodology (e.g., primer selection, volume filtered, filter pore size or size fractionation, and number of samples), as well as study site (e.g., Atlantic or Pacific) and seasonal timing of sampling linked to varied environmental conditions that shape the community (Govindarajan et al., 2023a). For example, our study was conducted in the productive Northwest Atlantic Slope Water region and consisted of bulk water sampling (no size fractionation) in March 2020, while the Blanco-Bercial et al. study (2022) was conducted monthly over the course of one year and the Giner et al. study (2020) used size-fractionated water samples (0.2-3 µm). Additionally, in our study, ASV accumulation curves for protists, invertebrates, and vertebrates suggested that the sampling scheme did not completely resolve the total biodiversity, which may also contribute to observed differences in diversity across studies.

Moreover, the chance encounter of a salp bloom during our sampling (see Methods) contributed to high relative abundance of salp sequences in surface waters (214) compared to other studies (Govindarajan et al., 2022; González et al., 2023; Matthews and Blanco-Bercial, 2023). Although we were unable to sample outside of the bloom to assess their impact on community composition, it is possible that diversity trends we observed may have been influenced by high salp densities. For example, previous studies suggest that salp blooms may impact biodiversity by substantially reducing phytoplankton biomass (Zeldis et al., 1995; Décima et al., 2023) and outcompeting other planktonic organisms for food (Paffenhöfer et al., 1995; Perissinotto and A. Pakhomov, 1998; Stukel et al., 2021).

Going forward, the maturation of multi-marker metabarcoding for deep sea studies will benefit from improved reference libraries for taxonomic assignments, increased spatial and temporal sampling, and standardization of sampling technologies, wet lab processes, and bioinformatics pipelines (McClenaghan et al., 2020; Patin and Goodwin, 2022; Kelly et al., 2023).

### 5.2 Eukaryotic community structure is depth-dependent

Eukaryotic communities were depth-dependent with differing community composition for protists, invertebrates, and vertebrates (Figure 3). Protist community composition was highly influenced by depth with distinct communities at each of our sampled depths and a notable shift in dominant taxa between epipelagic and mesopelagic depth zones. Invertebrate communities were also depth-specific, although there was more overlap across the communities with samples clustering into three depth bands for the epipelagic (< 200 m), upper mesopelagic (300-500 m), and lower mesopelagic (> 500 m). Vertebrate communities were less structured with depth with samples at 500 m exhibiting a significant difference between epipelagic and lower mesopelagic samples. The diminished depth clustering coincides with greater organismal mobility from single-celled protists to vertebrates, which may reflect a reduced depth constraint as swimming capacity increases.

Depth-dependency of eukaryotic communities, especially for protists, has been established previously, with a distinct transition in community composition between the sunlit surface and the deep (Ollison et al., 2021; Blanco-Bercial et al., 2022; Yeh and Fuhrman, 2022; Sun et al., 2023; Anderson et al., 2024). For protists, a high relative abundance of photosynthetic taxa including stramenopiles and chlorophytes transitions into dominance by alveolates, especially Syndiniales and radiolarians.

Similar to other metazoan studies, we found high relative abundance of calanoid copepods (Order Calanoida) in the water column, as well as a subsurface abundance of Siphonophorae and Ctenophora (e.g., comb jellies) (Laroche et al., 2020b; Govindarajan et al., 2021). For vertebrates, we detected high relative abundance of mesopelagic fish from families Myctophidae and Gonostomatidae, which were also present in other metabarcoding studies (Canals et al., 2021; Govindarajan et al., 2023b; Dan et al., 2024). In epipelagic waters, additional taxa occurring in high relative abundance included the families Serrivomeridae (sawtooth eels), Molidae (molas), and Alepocephalidae (slickheads).

We did not detect strong signals of diel vertical migrators in our 18S and 12S datasets (Figure 3). The 18S V9 biomarker used in this study amplifies many protistan and invertebrate taxa, which can dilute the signal of migrating taxa. In a previous study with higher diel sampling resolution, DVM was also not detected when using the 18S biomarker (Govindarajan et al., 2021). For fishes, previous studies detected higher deep-sea species richness at night in surface waters (Canals et al., 2021; Govindarajan et al., 2023b). However, we noted mainly lower species richness at surface depths (Supplementary Table S11). The lack of strong DVM signals in our datasets is likely due to a combination of factors including the sensitivity of eDNA approaches, low diel sampling resolution with only three CTD casts, and the biomarker choice. Moreover, while eDNA metabarcoding can capture a robust snapshot of the whole community biodiversity, it may not accurately reflect the active community as eDNA in the water column originates from both dead and living organisms and encompass whole cells and genetic material that is sloughed off or excreted from moving animals (Allan et al., 2021b; Rodriguez-Ezpeleta et al., 2021). For example, salps are known diel vertical migrators (Wiebe et al., 1979; Madin et al., 2006) and we observed in situ DVM (Supplementary Figure S2). However, eDNA depth profiles were not as clear with higher relative abundance in deeper waters during the night and day (Supplementary Figure S24), which may be a result of eDNA sourced directly from salps or from sinking fecal pellets (Govindarajan et al., 2021).

To probe potential signals of DVM, we instead focused our analyses on the presence of unique, shared, and cosmopolitan ASVs across depth (Supplementary Figure S11). The detection of unique ASVs could indicate niche specialization to different environmental conditions within a given taxonomic group (Ollison et al., 2021; Anderson et al., 2024). Unexpectedly, we found unique ASVs for diatoms and chlorophytes in deep waters. Rather than niche specialization of these photosynthetic taxa to aphotic conditions, we hypothesize that these sequences indicate export of primary producers out of the euphotic zone as sinking particles (Amacher et al., 2013; Durkin et al., 2016, 2021, 2022; Mestre et al., 2018; Baumas and Bizic, 2024; Bodel et al., 2025; Kramer et al., 2025).

Animals shed DNA as they move through the water column, and we hypothesize that unique ASVs represent cohorts of animals that are non-migratory while shared and cosmopolitan ASVs reflect migratory cohorts. However, another possible explanation for the high occurrence of unique invertebrate and vertebrate ASVs may be due to under-sampling of metazoan biodiversity, which tend to be patchily distributed in the water column. Accumulation curves for ASV richness across our sample set suggest that additional sampling may be necessary to recover the true community richness for metazoans, especially for fish (Supplementary Figure S25). Environmental factors that influence eDNA shedding, persistence, and transport in combination with low-volume (∼1-5 L) sampling efforts impact eDNA recovery for metazoan detection (Govindarajan et al., 2022, 2023a). While these volumes may be adequate for microbial studies, they likely underestimate metazoan biodiversity in deep sea ecosystems (Govindarajan et al., 2022), which can lead to a skewed perspective of their vertical and horizontal distribution. However, unique and cosmopolitan ASVs can be used to probe DVM behaviors, which may be useful to better understand the ecology of these otherwise elusive taxa that warrant further study.

We detected invertebrate and vertebrate ASVs that appeared at multiple sampling depths (shared and cosmopolitan ASVs), which included representatives from known migratory invertebrate taxa including copepods, cnidarians, and tunicates (Bianchi and Mislan, 2016; Romero-Romero et al., 2019; Easson et al., 2020), as well as mesopelagic fish including lanternfish and bristlemouths. Most of these fish are considered migratory except for *Cyclothone microdon* (bristlemouth) and *Scopeloberyx opisthopterus* (bigscale), which are lower mesopelagic and bathypelagic dwelling (Sutton, 2013; Govindarajan et al., 2023b; Richards et al., 2023). However, our study detected one ASV each of *C. microdon* (15 ASVs total) and *S. opisthopterus* (97 ASVs total) with a relatively high abundance in epipelagic waters, especially during the day. While studies have indicated that juvenile stages of mesopelagic fish may occur at shallower depths (Olivar and Beckley, 2022), it remains unclear whether the driving mechanism is ontogenetic migration or other drivers such as niche expansion that require further study (Sutton et al., 2008; Cook et al., 2013; Granata et al., 2023).

### 5.3 Protists increase ecosystem complexity and stability

Diversity (nodes or ASVs), connectivity (edges or interactions) and edge density (edge-to-node ratio) are core determinants of network complexity (Landi et al., 2018; Espinoza et al., 2020). Increasing biological diversity and their interactions increase ecosystem complexity, which often correlates with enhanced stability or resilience (McCann, 2000; Dunne et al., 2002; Carpentier et al., 2021; Nauta and De Domenico, 2024). Our ecological network analyses from multi-marker metabarcoding of the eukaryotic community showed that protistan taxa play a key role in mediating trophic interactions and ecosystem function. While metazoan networks were fragmented, the addition of protistan ASVs increased diversity and interactions (Supplementary Table S7), producing fully connected networks at all depths with protist nodes facilitating interactions between previously unconnected metazoan nodes (Figure 5). Increased protistan diversity may stabilize connectivity by enhancing the likelihood that biological communities are resilient to environmental fluctuations and can compensate the loss of functionally important taxa (functional redundancy) (Naeem and Li, 1997; Naeem, 1998; McCann, 2000).

Alternatively, our fragmented ecological networks may reflect the more heterogenous distributions of metazoan taxa that reduce co-occurrences and interactions, especially in the vast and deep pelagic ocean where food is scarce (Robison, 2004; Ramirez-Llodra et al., 2010). Fragmented networks may also reflect sampling biases associated with low-volume samples, which may not provide a comprehensive snapshot of metazoan diversity and distribution compared to more easily sampled protistan taxa (Anderson and Thompson, 2022; Govindarajan et al., 2022). Significant knowledge gaps on metazoan eDNA further complicate our interpretation of metazoan eDNA signatures due in part to unknown rates of DNA shedding across different taxa and the varying impacts of environmental parameters on eDNA persistence (Allan et al., 2021a, 2021b; McCartin et al., 2022; Govindarajan et al., 2023a). Nevertheless, molecular tools like multi-marker metabarcoding and increasing availability of large-volume autonomous samplers are revolutionizing biodiversity research and provide important multi-trophic insights that are otherwise difficult to obtain, especially in the deep sea (Govindarajan et al., 2023a; Dan et al., 2024; Haddock and Choy, 2024; Yang et al., 2024).

### 5.4 Protists are key mediators of ecological interactions

Positive edges indicate ASVs that tend to exhibit similar co-variation patterns across samples, whereas negative edges indicate opposing patterns. While signed associations do not imply direct interactions, they provide a useful framework for generating hypotheses about facilitative or antagonistic relationships that can be tested in future studies. Consistent with other network studies, we interpret these associations as potentially reflecting shared or divergent ecological responses. For example, positive interactions may indicate putative mutualistic or commensal relationships while negative interactions suggest antagonistic relationships including competition, predation, and parasitism (Xiao et al., 2017; Landi et al., 2018; Anderson et al., 2024; Deutschmann et al., 2024). The balance of positive and negative interactions has been shown to shape community interactions and ecosystem processes and in turn, ecological stability and resilience to environmental change (Travis et al., 2005; Losapio et al., 2021). In our networks, there were more positive interactions than negative interactions across all depth zones (Supplementary Table S8), indicating an important role for mutualistic or facilitative relationships in marine communities (Stachowicz, 2001; Deutschmann et al., 2024). The proportion of negative interactions increased with depth, which may suggest that competition or predation dynamics increases in importance in the relatively resource-scarce mesopelagic.

Positive networks were also more robust to node loss than negative networks and for both network types, mesopelagic networks were more robust (Supplementary Figure S16). This may be a result of the higher complexity found in the positive networks that can enhance network stability. Moreover, positive networks had higher modularity, suggesting increased compartmentalization of ASVs with potential niche partitioning and functional specialists (Supplementary Table S8), which may also have a moderately stabilizing effect on networks. This trend did not apply to depth-specific networks where positive and negative networks in the epipelagic had higher modularity but lower robustness than mesopelagic networks. In our study, decreased complexity in the epipelagic relative to mesopelagic depths is likely a stronger driver of network stability. These dynamics suggest that negative-type interactions (e.g., key predator-prey relationships), particularly in the epipelagic, may be more vulnerable to fragmentation or loss of keystone taxa.

Top-ranking nodes indicating keystone taxa are highly connected ASVs that are important facilitators of resource transfer (i.e., carbon and energy) throughout a network. A majority of the top-ranking nodes in our study were protists (Supplementary Figures S18-S19), which included a high number of ASVs belonging to the marine alveolate Syndiniales, suggesting an important role for parasitism in the marine environment (Guillou et al., 2008; de Vargas et al., 2015; Anderson et al., 2024). Copepod ASVs as well as various gelatinous taxa (cnidarians, ctenophores, and salps), mesopelagic fish, and top predators (such as Risso’s dolphin and common dolphinfish) were also identified as keystone taxa (Supplementary Figures S18-19). Our findings highlight protists as keystone taxa while also underscoring the importance of maintaining biodiversity across trophic levels and functional groups to sustain ecosystem function and stability.

We identified known and previously unobserved trophic interactions in our simplified food web analysis that featured protist associations with both invertebrates and vertebrates (Figure 6). Protist-based associations were more common than invertebrate-vertebrate associations, which were likely removed from consideration due to the low abundance and prevalence of metazoan ASVs, especially for fish. Protist-driven associations likely play a key role in carbon export out of the surface ocean and through deep sea food webs. For example, sinking particles and marine snow harbor microbial communities comprised of senescent primary producers and diverse microbial communities. Thus, our finding of protist-dominated interactions, especially in mesopelagic networks, aligns with the consensus that marine snow is an important food source in the deep sea (Dilling and Brzezinski, 2004; Choy et al., 2015; Cawley et al., 2021). Additionally, these putative interactions likely do not reflect direct feeding relationships between protists and metazoans, but may highlight indirect relationships between microorganisms and metazoans linked by plankton-particle dynamics that contribute to overall ecosystem function and warrant further study (Djurhuus et al., 2020; Holman et al., 2021; Doherty et al., 2025).

## 6. Conclusions

Mesopelagic studies that integrate both microorganisms and metazoans are rare, yet growing anthropogenic pressures underscore the need for more holistic research on this critical ocean realm. Our study applied a multi-marker metabarcoding approach to survey mesopelagic biodiversity in the Northwest Atlantic Slope Water region and construct ecological networks to examine mesopelagic diversity, organismal interactions, and ecosystem stability. Our findings revealed depth-specific structuring of the eukaryotic community, spanning protists, invertebrates, and vertebrates. We did not detect clear DVM patterns in our overall dataset, likely due to limited diel sampling. Assuming resource availability and feasibility, future sampling efforts should span across multiple day-night cycles to increase diel resolution. In addition, since eDNA originates from both living and dead organisms, incorporating environmental RNA as a more accurate approach for tracking DVM activity should be considered since RNA is only shed by living organisms (Cristescu, 2019; Yates et al., 2021).

Notably, ecological networks that included all eukaryotic groups were more robust than metazoan-only networks, with the incorporation of protist ASVs enhancing network complexity and stability. The higher diversity of protistan ASVs likely conferred resilience to taxa loss across mesopelagic networks compared to the epipelagic network. Additionally, we found that negative interactions were more vulnerable to taxa loss than positive interactions, with implications for carbon and energy transfer within food webs and deep-sea carbon sequestration. While mesopelagic networks appear more structurally stable, it is crucial to recognize that surface and deep-sea ecosystems are intrinsically linked through the biological carbon pump and vertically migrating organisms. Therefore, a more complete understanding of biodiversity, species interactions, and their impact on ecosystem function across spatial, temporal, and taxonomic scales is essential for effective ocean governance of the deep pelagic ocean.

Future work incorporating a cross-domain lens, inclusive of bacteria and archaea, and biogeochemical processes such as carbon flux (Guidi et al., 2016; Nguyen et al., 2022; McMonagle et al., 2024) will help us better understand the role of biodiversity and their interactions in facilitating carbon cycling and other critical ecosystem services.

## 7. Acknowledgments

We are grateful for everyone involved in the collection of environmental DNA samples including Rene Francolini and the crew of the R/V Armstrong. We also thank Erin Frates for laboratory processing of the samples. This research is part of the Woods Hole Oceanographic Institution’s Ocean Twilight Zone Project, supported by funding as a part of The Audacious Project, a collaborative endeavor housed at TED and also funded in part by the National Oceanic and Atmospheric Administration’s (NOAA’s) Ocean Exploration Cooperative Institute under award NA19OAR4320072 to the University of Rhode Island (subaward number is 0007525/102212019 to WHOI).

## 9. Data accessibility statement

All relevant data, metadata, and code used in this study are available openly available on GitHub at https://github.com/yang-nina/OTZ-AR43-Metabarcoding. Raw sequence reads are deposited in the SRA (BioProject accession PRJNA1454214).

## 10. Benefit-sharing statement

Benefits of this research are shared as all relevant data, results, and code necessary to reproduce this study.

