## Supplemental Materials for "Eukaryotic biodiversity and ecological networks from the surface to the mesopelagic in the Northwest Atlantic Slope Water"

### Table of Contents

1. Supplementary Methods
2. Supplementary References
3. Supplementary Discussion
4. Supplementary Figures S1 to S25
5. Supplementary Tables S1 to S11

### 1. Supplementary Methods

Acoustic methods

The R/V Armstrong is equipped with a broadband Simrad EK80 split beam echosounder system (Kongsberg Maritime) with 5 acoustic channels (200, 120, 70, 38, and 18 kHz transducers). The transceiver was configured with a K-Sync module to transmit each frequency in sequence. Calibration parameters were obtained for each acoustic frequency by the standard sphere method prior to cruise work (March 2019), except for the 18 kHz where the factory calibration was used (De Robertis and Higginbottom, 2007). There were several sources of noise in the data, including from interference with other acoustic systems, bubble-dropouts, and changes to the noise floor. The acoustic data was first processed with a custom MATLAB (MATHWORKS, Inc.) library and Volume Scattering Strength (Sv) was calculated following (Andersen et al., 2024) in 1-m range bins. Bubble dropouts were removed by excluding pings with Sv values below -90 dB continuously for > 20 m range. Pings that were affected by strong acoustic interference (Sv > -40 dB) over long ranges were also removed. Pings affected by an elevated noise floor were removed by setting an upper threshold on the total Mean Volume Backscattering Strength (MVBS) calculated in the relatively low scattering region between 900 and 1000 m. Sv data were averaged in 1-minute bins following the noise mitigation steps as displayed in the echogram. Median Sv profiles were then calculated per frequency across the echogram duration.

Profiles for the environmental parameters (temperature, salinity, oxygen, chlorophyll-a) were calculated in 1-m depth bins averaged across the dataset.

Imaging Methods

Salps were enumerated from high resolution shadowgraph image data analyzed with a machine learning model trained for the detection of zooplankton and micronekton taxa (YOLOV8). On the hold out image dataset used for model evaluation, an F1 score of 0.81 was achieved for the salp category at the confidence score used for processing (0.7). Salp concentrations with 95% confidence interval were calculated from 25-m depth-binned count data and the corresponding volume of water imaged.

Decontamination of metabarcoding datasets

Additional clean-up was conducted manually to remove unwanted ASVs. Following the use of decontam in R, the protist dataset had 9737 ASVs. Further processing removed ASVs that were in the domain Eukaryota:nucl, and ASVs that were not assigned at the supergroup level, resulting in a total of 7127 ASVs retained. For the invertebrate dataset, decontam resulted in a total of 480 ASVs. Additional processing removed ASVs that were unassigned or assigned as Archaea at the kingdom level, as well as ASVs assigned as Vertebrata, class Arachnida, and class Insecta, for a final dataset comprised of 404 ASVs prior to rarefaction. For the vertebrate dataset, decontam resulted in 859 ASVs. Further processing removed 132 contaminant ASVs that were assigned with a custom BLAST search as terrestrial mammals including *Canis lupus* (gray wolf), *Gallus gallus* (chicken), and *Bos taurus* (cattle). Additionally, one ASV assigned as genus *Fundulus* (freshwater fish) and one ASV assigned as species *Centropyge acanthops* (popular aquarium fish) were removed. This resulted in a final dataset of 727 ASVs prior to rarefaction.

### 3. Supplementary Discussion

We pooled samples across CTD casts that were sampled during the day and night, which may dilute signals of diel vertical migration. We conducted PERMANOVA analyses to look at the comparison between CTD casts at each depth and found no clear signs of DVM despite some statistically significant pairwise comparisons. These significant results likely reflect community variation across replicates and CTD casts as well as other sampling and methodological trade-offs. For example, there is fine-scale variability in biomass distributions with depth and at different stations, but the sampling depths were pre-set and targeted one scattering layer at ~500m. This could lead to varied relative abundance profiles across stations. As a result, pooling our samples is the appropriate approach for our work.

**Protists:** We found statistical significance ($p < 0.05$) between CTD casts at each depth (Casts 001, 002, and 007 are different from each other). This result does not reflect clear DVM although some protists are known migrators. Instead, we interpret this to reflect natural variation in the community and/or methodological artifacts. While 5L of water is considered low-volume for metazoans, it is a relatively large volume for microbes where sampling is often 0.5L-1L. Here, the "high" volume captures a very high-resolution perspective of the community where small differences in relative abundance can skew the statistics. Relative abundance profiles for each CTD cast showed similar depth profiles for protists (Supplementary Figure S8).

**Invertebrates**: We found statistical significance ($p < 0.05$) between Cast 002 and Cast 007 at 10m, which may reflect some signals of DVM as salps are abundant in Cast 007 compared to Cast 002. However, the signal is not clear given the lack of significance between Cast 001 and Cast 002. In fact, higher salp relative abundance in deeper waters at night (Cast 001 and Cast 007) were observed than during the day (Cast 002). This may be due to collecting DNA from salp fecal pellets at depth or a residual eDNA signature from the previous day. It may also be due to the use or relative abundances instead of absolute abundance, which is a limitation of using eDNA.

### 4. Supplementary Figures

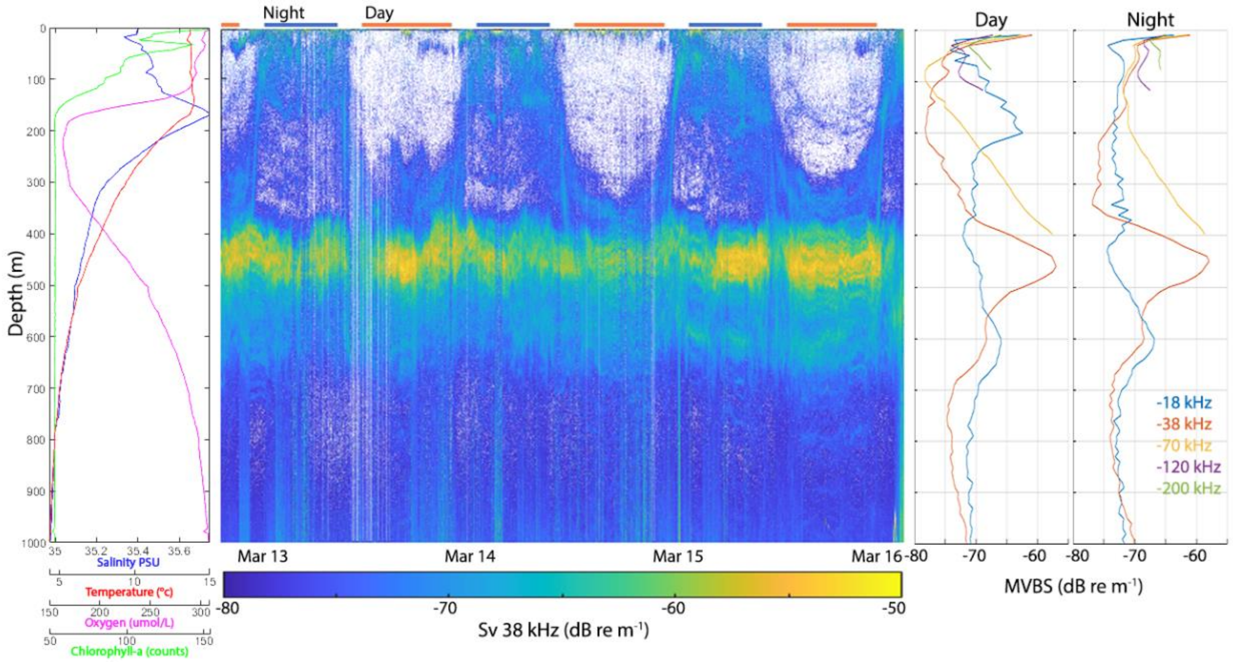


**Supplementary Figure S1.** Observations from AR43 in the Northwest Atlantic Slope Water region showing vertical profiles of environmental properties and a multi-day 38 kHz echogram with a prominent deep acoustic scattering layer in the depth range ~400-500 m. See Supplementary Methods for additional details.

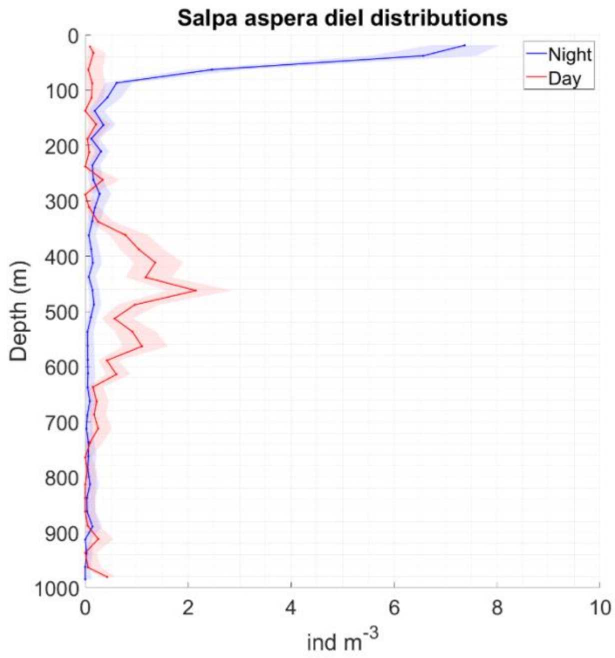


**Supplementary Figure S2.** Salp density estimates derived from high resolution shadowgraph image data. Abundance estimates in 25-m depth bins are shown for pooled daytime (red) and nighttime (blue) vertical profiles. Shaded area shows the 95% confidence interval.

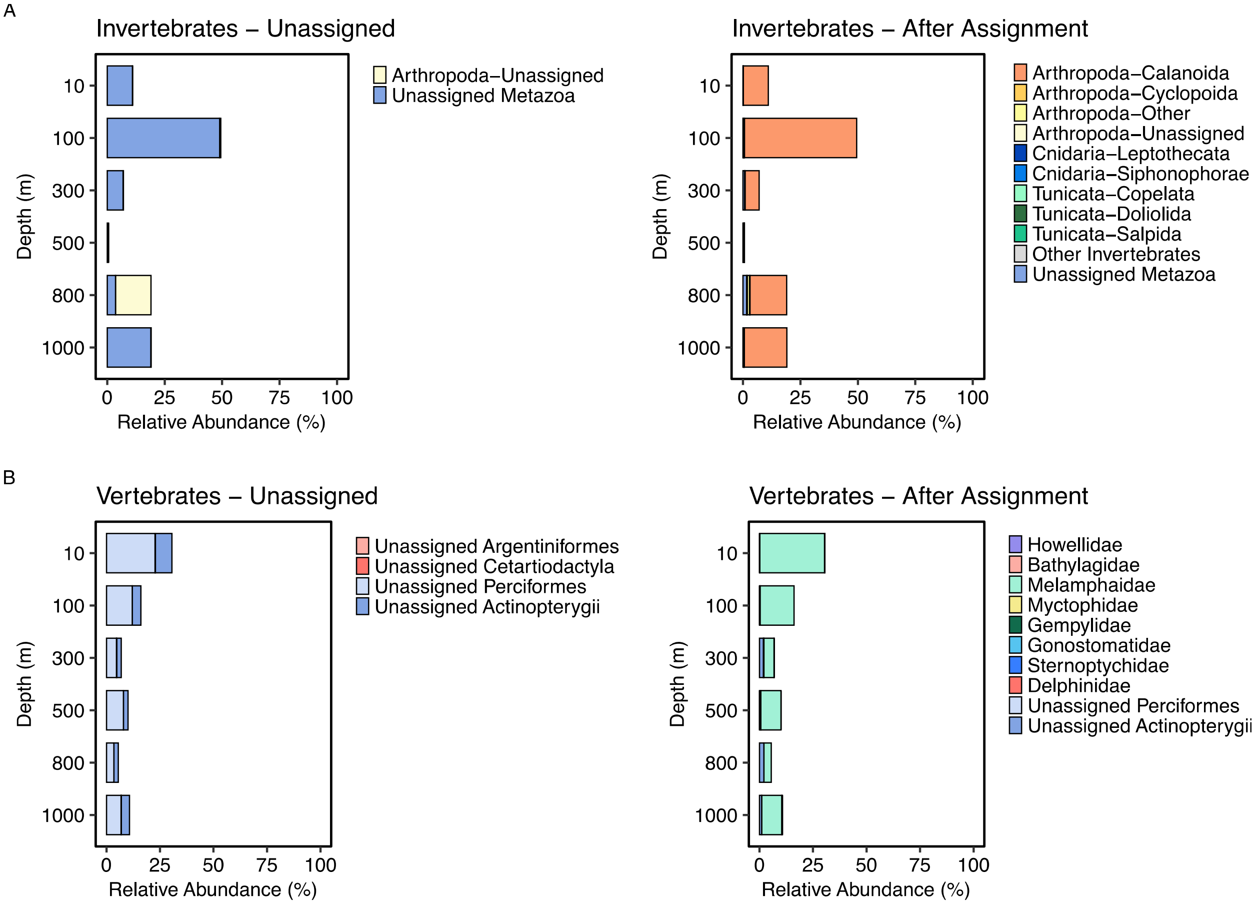


**Supplementary Figure S3.** A) Invertebrate and B) vertebrate depth profiles of percent relative abundance for unassigned sequences following taxonomic classification with a classifier-based approach in QIIME 2 (left) and after manual curation of unassigned sequences with the BLAST search function in Geneious Prime 2025.0.3 (right). Samples across the three CTD casts were aggregated to calculate relative abundance across the entire dataset. For invertebrates, taxonomic groups (Phylum and Order) are colored and listed in alphabetical order. "Other Invertebrates" includes the following phyla: Annelida, Mollusca, Nematoda, and Rotifera. "Other Arthropoda" includes the orders Amphipoda, Decapoda, Harpacticoida, Mormonilloida, and Scalpellomorpha. Vertebrate families were grouped at the Order level: Howellidae (Acropomatiformes); Bathylagidae (Argentiniformes); Melamphaidae (Beryciformes); Myctophidae (Myctophiformes); Gempylidae (Scombriformes); Gonostomatidae, Sternoptychidae (Stomiiformes); and Delphinidae (Cetartiodactyla). Note: Unassigned taxa are listed as their lowest assigned taxonomic rank. Invertebrates were considered unassigned when Phylum and/or Order were unresolved. Vertebrates were considered unassigned when Family was unresolved.

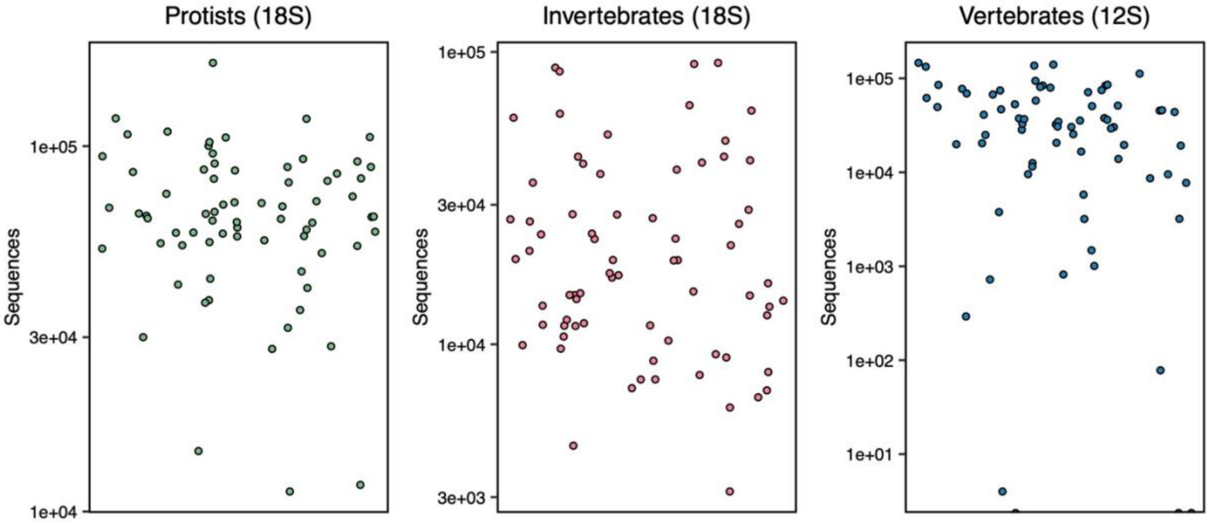


**Supplementary Figure S4.** Sampling coverage plots were used to select a sequencing threshold to remove low read-depth outliers. Each sample is represented by a point with a randomized position to avoid overlap on the x-axis. The y-axis shows the sequencing depth as a $\log_{10}$ scale.

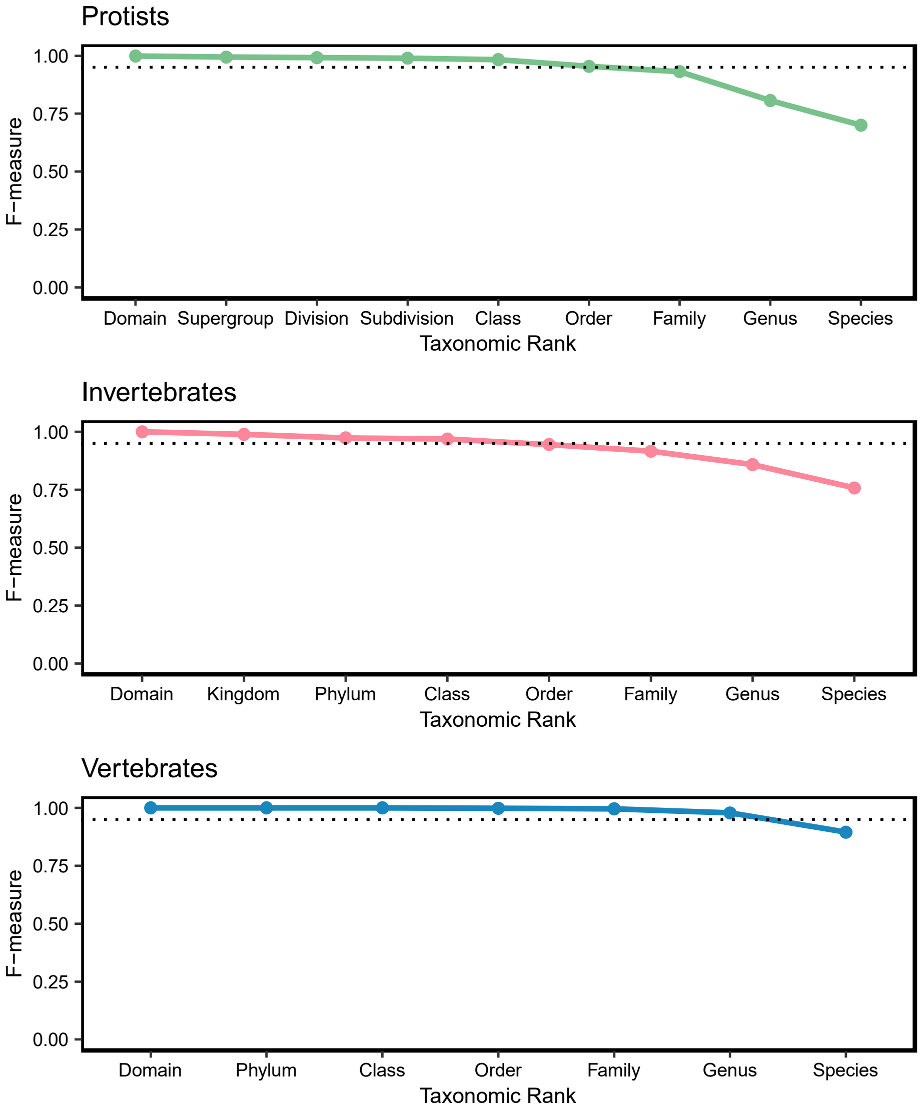


**Supplementary Figure S5.** Naïve Bayes classifier performance was evaluated for the protist (18S V9, PR2), invertebrate (18S V9, SILVA), and vertebrate (12S MiFish, MetaZooGene with custom database from Govindarjan et al., 2023) classifiers. The x-axis shows taxonomic rank, and y-axis is the F-measure, a metric that reflects a balance between precision and recall of the classifiers as evaluated with the RESCRIPt function evaluate-fit-classifier in QIIME 2. The dotted line represents a cutoff of 0.95 as a threshold for high performance.

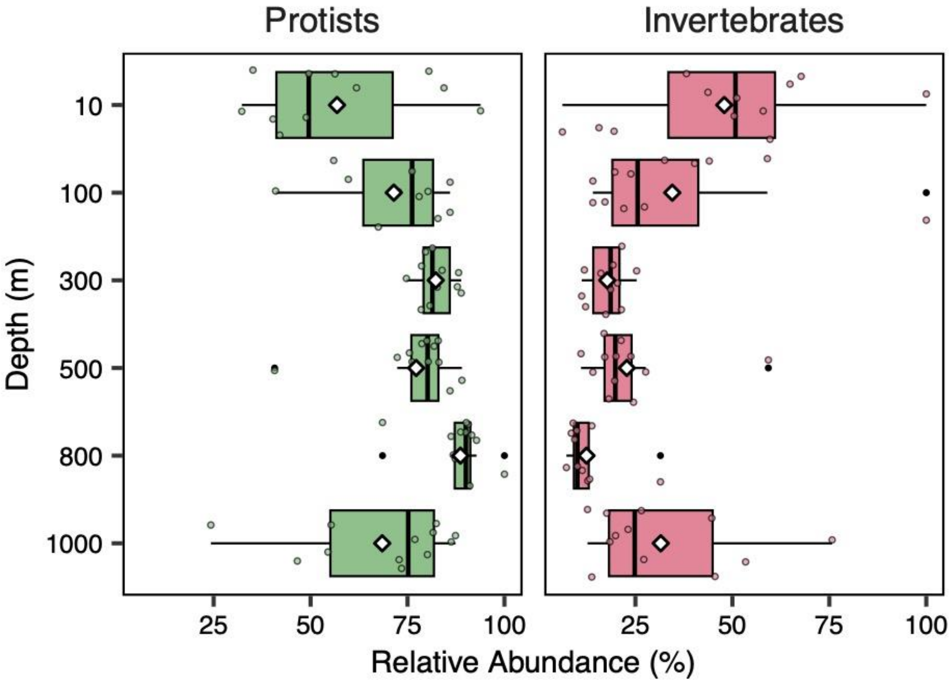


**Supplementary Figure S6.** Proportion of sequencing reads for protists (n=69) and invertebrates (n=70) from 18S V9 metabarcoding. Boxplots show the variation in the proportion of reads within a sample between depths divided into quartiles with outlier values indicated by black points. A white diamond indicates the mean value at each depth.

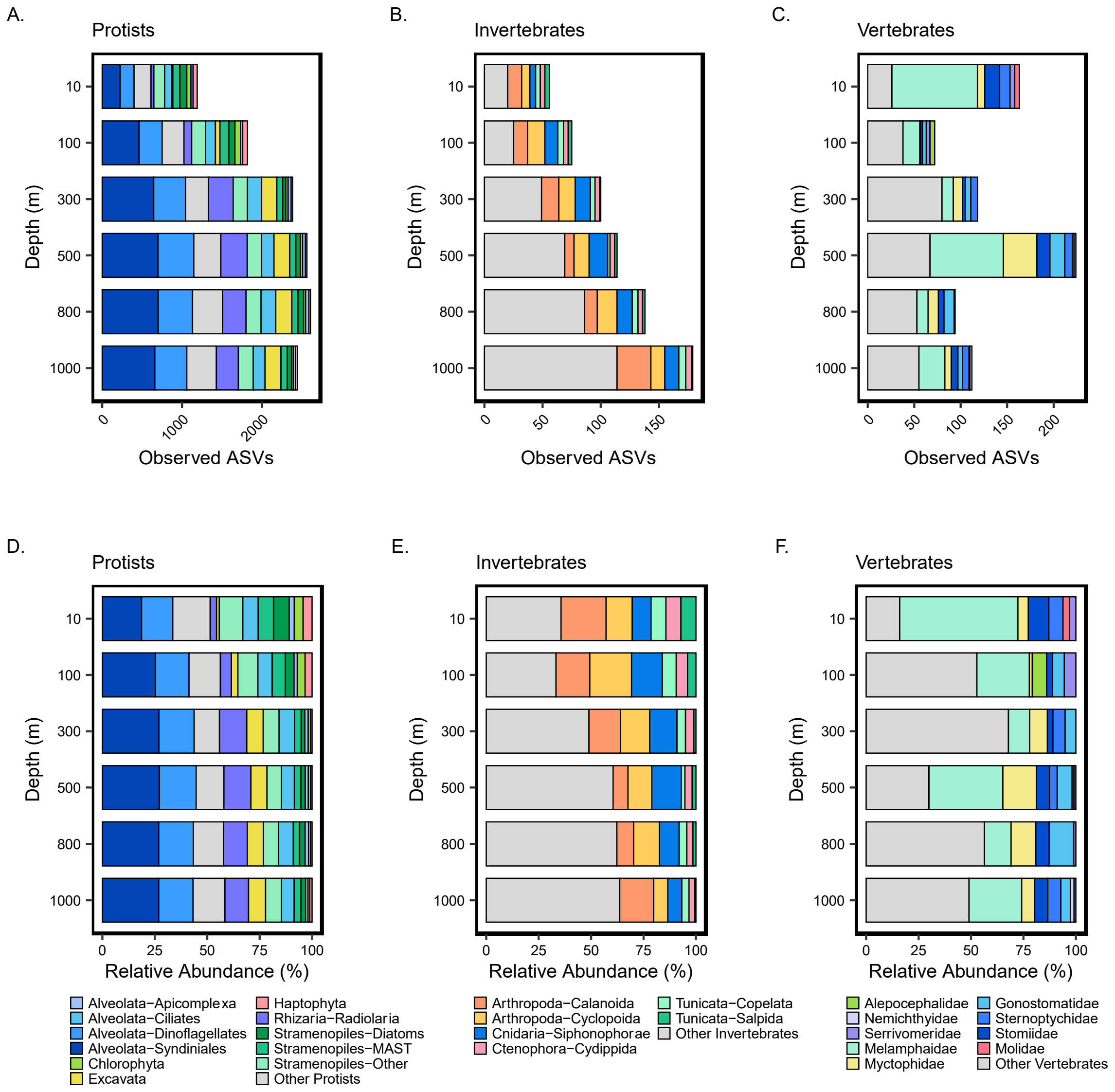


**Supplementary Figure S7.** Depth profiles of genetic diversity as the number of unique observed ASVs and the relative proportion (%) of ASVs at each depth for A & D) protists, B & E) invertebrates, and C & F) vertebrates. Bar plots show taxonomic groups in the order of decreasing proportion (%). In the legend, taxa within protists (Supergroup and/or Division or Subdivision) and invertebrates (Phylum and Order) are colored and listed in alphabetical order. Vertebrate families were grouped at the Order level in alphabetical order: Alepocephalidae (Alepocephaliformes); Nemichthyidae, Serrivomeridae (Anguilliformes); Melamphidae (Beryciformes); Myctophidae (Myctophiformes); Gonostomatidae, Sternoptychidae, Stomiidae (Stomiiformes); and Molidae (Tetraodontiformes). For each group, only taxa with overall relative abundance > 1% are displayed, with all other taxa combined in “Other”. For a full list of “Other” taxa, see Supplementary Tables S1-S3.

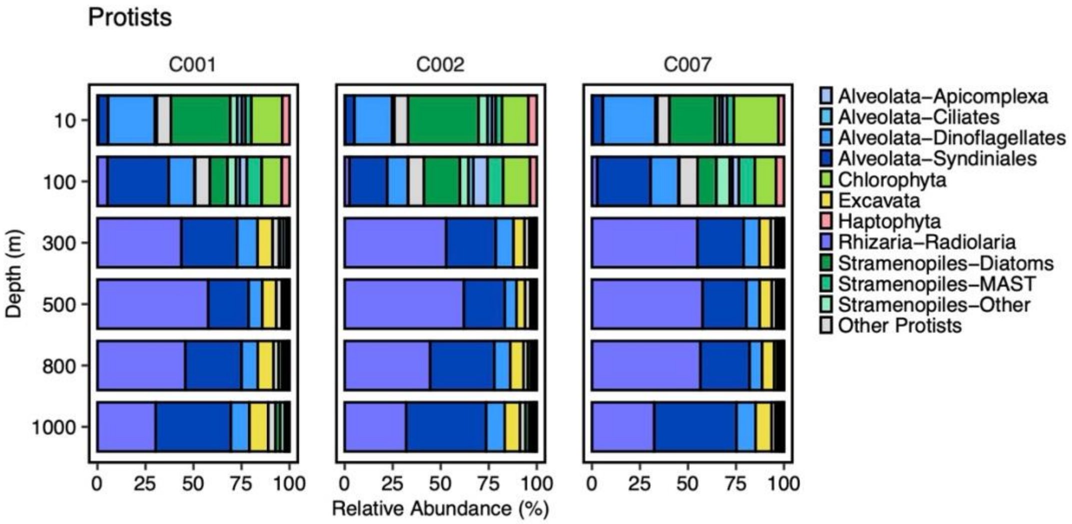


**Supplementary Figure S8.** Depth profiles of relative read abundance (%) for protists shown for each deployment. For each taxonomic group, only taxa with overall relative abundance > 1% are displayed. Cast 001 and C007 were collected at night while Cast 002 was collected during the day.

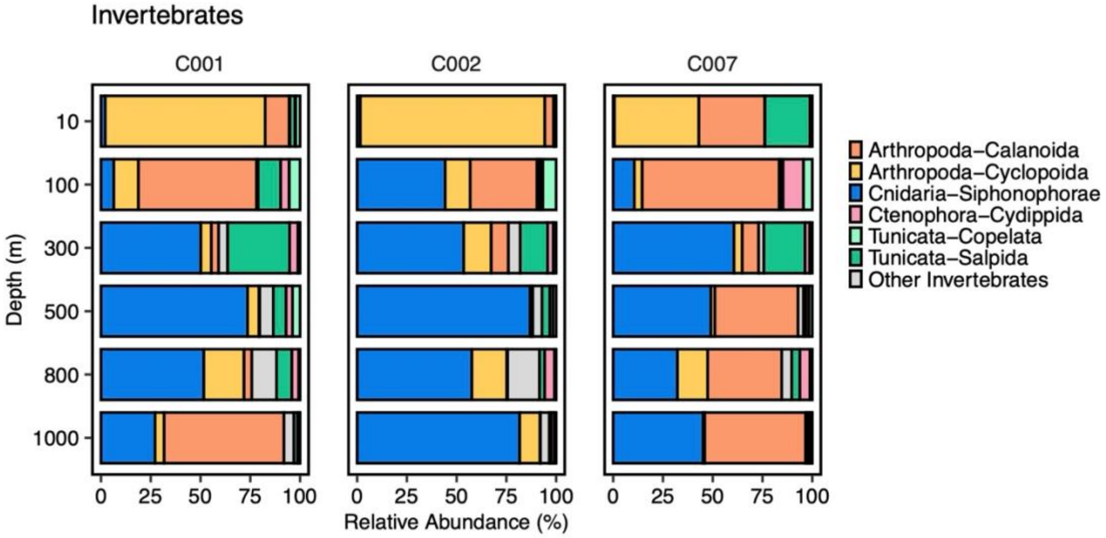


**Supplementary Figure S9.** Depth profiles of relative read abundance (%) for invertebrates shown for each deployment. For each taxonomic group, only taxa with overall relative abundance > 1% are displayed. Cast 001 and C007 were collected at night while Cast 002 was collected during the day.

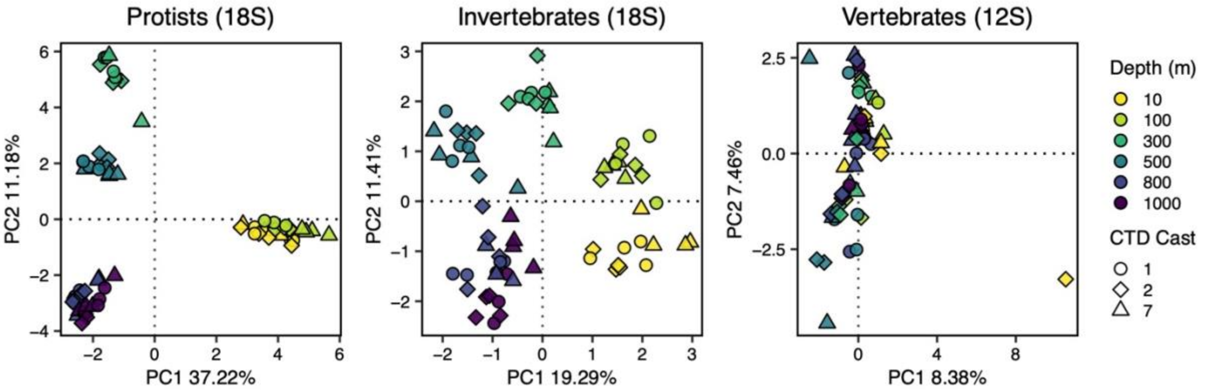


**Supplementary Figure S10**. Principal component analysis (PCA) of center-log ratio-transformed reads for protists (n=69), invertebrates (n=70), and vertebrates (n=61). Each point represents one sample. The shape and color of each point indicates the CTD cast and the sampling depth, respectively, of each sample. CTD cast here can also reflect diel sampling with casts 1 and 7 sampled at midnight and cast 2 sampled during mid-day. In the vertebrate PCA, the outlier at 10 m (PC1 = ~10) is retained.

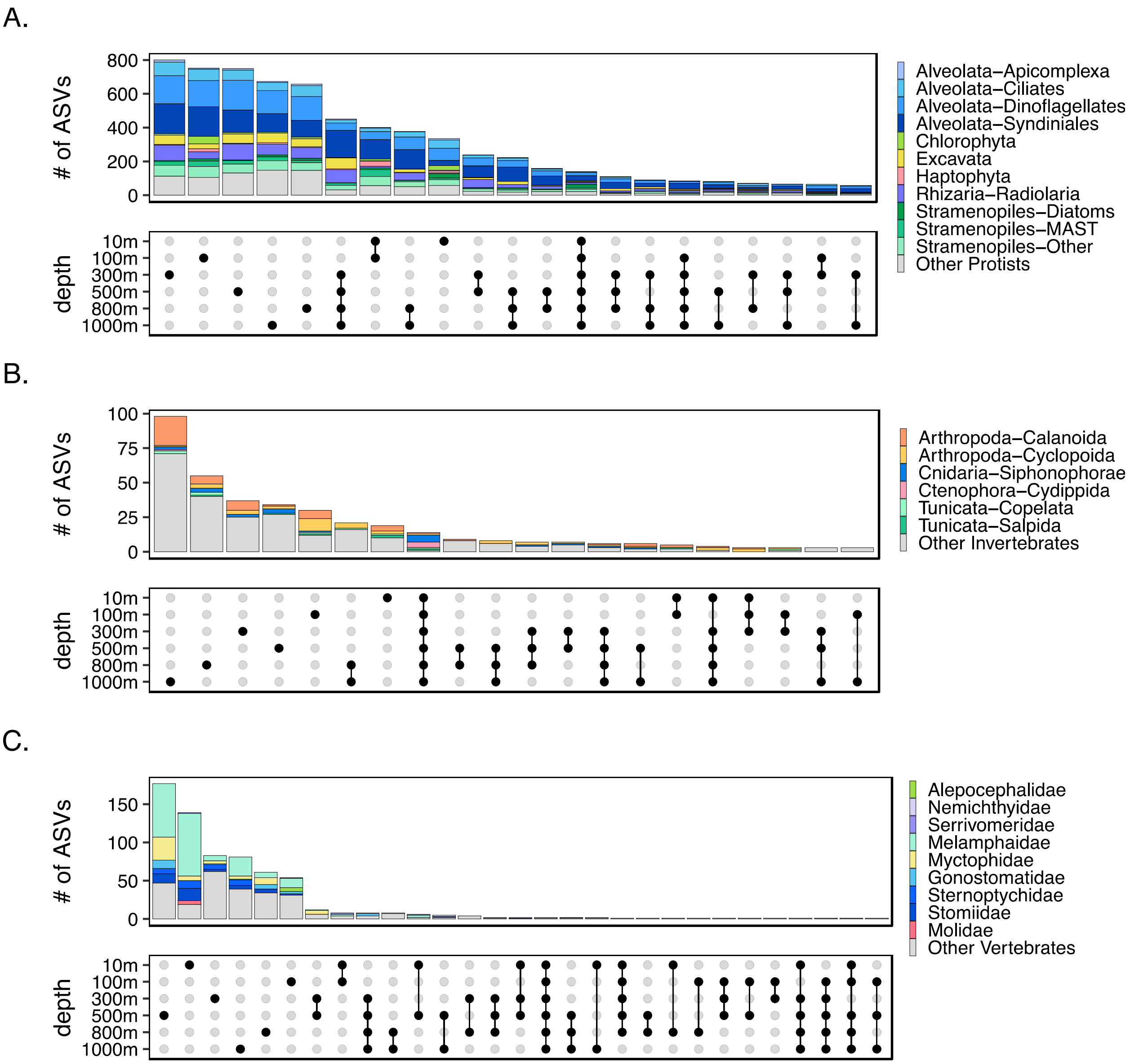


**Supplementary Figure S11.** UpSet plots for A) protists, B) invertebrates, and C) vertebrates show how ASVs overlap or intersect across depth. In the dot grid or intersection matrix (bottom), each row represents a depth, and each column represents an intersection. Single filled dots in black indicate unique ASVs while connected dots in black indicate shared ASVs across multiple depths. The intersection size bar plot (top), shows how many ASVs are in each combination of depths. The sets of ASVs are colored based on key taxonomic groups. In the legend, taxonomic groups within protists (Supergroup and/or Division or Subdivision) and invertebrates (Phylum and Order) are colored and listed in alphabetical order. Vertebrate families were grouped at the Order level in alphabetical order: Alepocephalidae (Alepocephaliformes); Nemichthyidae, Serrivomeridae (Anguilliformes); Melamphidae (Beryciformes); Myctophidae (Myctophiformes); Gonostomatidae, Sternoptychidae, Stomiidae (Stomiiformes); and Molidae (Tetraodontiformes). For each group, only taxa with overall relative abundance > 1% are displayed, with all other taxa combined in “Other”. For a full list of “Other” taxa, see Supplementary Tables S1-S3.

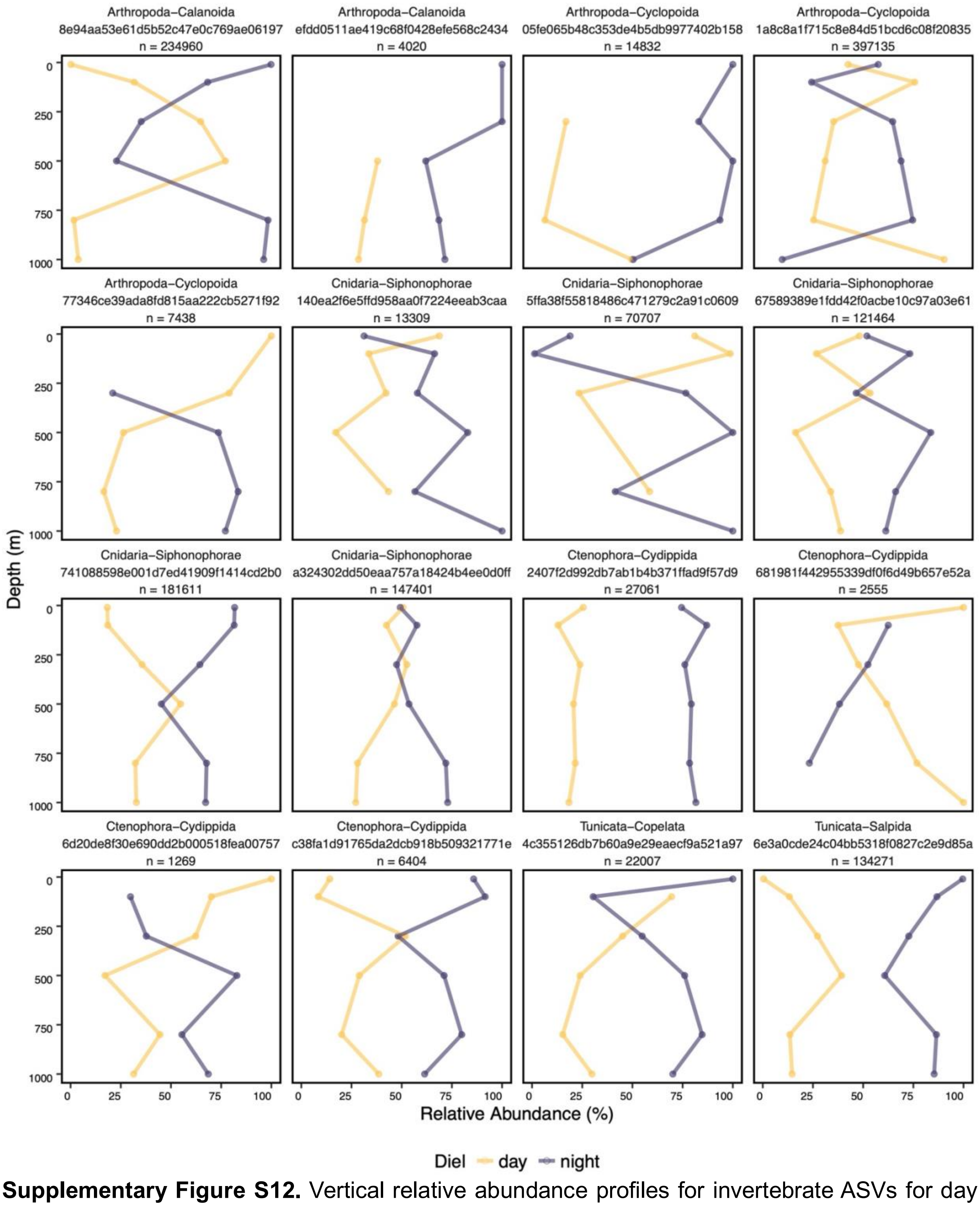


**Supplementary Figure S12.** Vertical relative abundance profiles for invertebrate ASVs for day (cast 2) and night (cast 1 and 7). The invertebrate ASV and taxonomic classification (Phylum-Order), as well as the total number of reads for the ASV, are shown in the facet header. Relative read abundance is shown on the x-axis and depth (m) is shown on the y-axis. Only ASVs that occurred at multiple depths spanning the water column were included.

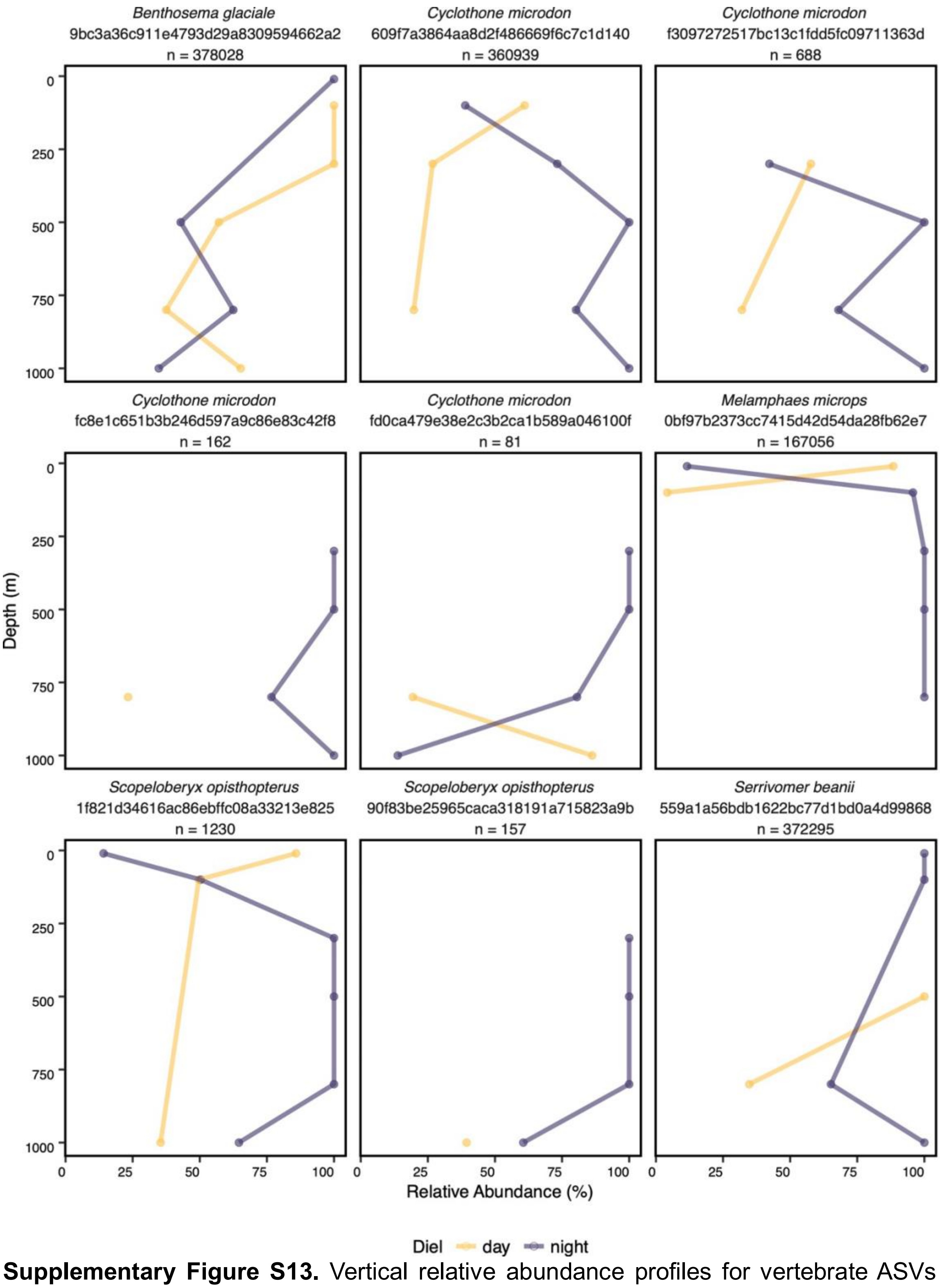


**Supplementary Figure S13.** Vertical relative abundance profiles for vertebrate ASVs for day (cast 2) and night (cast 1 and 7). The vertebrate ASV and species name, as well as the total number of reads for the ASV, are shown in the facet header. Relative read abundance is shown on the x-axis and depth (m) is shown on the y-axis. Only ASVs that occurred at multiple spanning the water column were included.

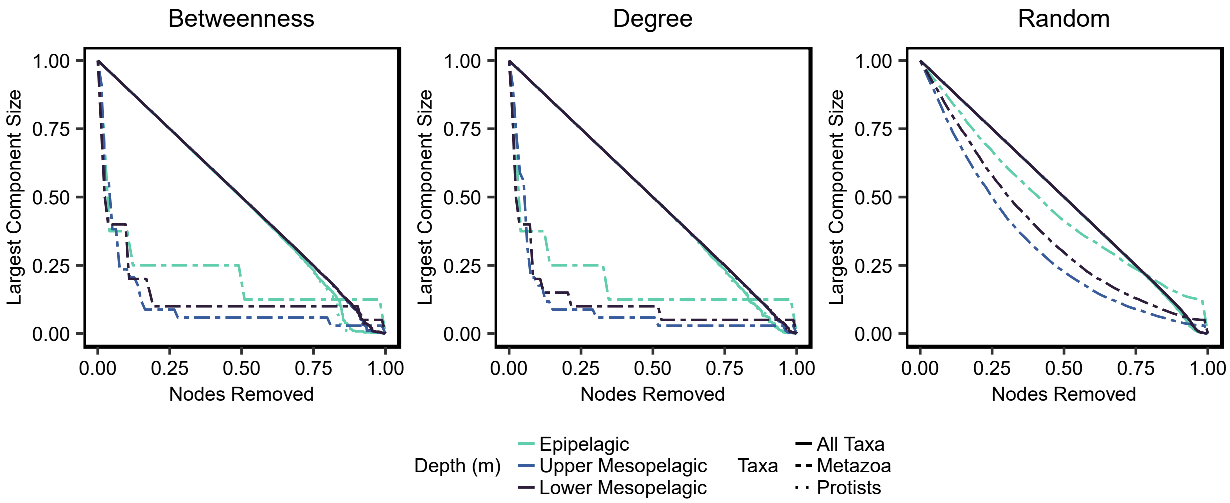


**Supplementary Figure S14.** Robustness curves for All Taxa, Metazoa-only, and Protists-Only networks at each depth zone. Attack robustness of each network was measured by sequentially removing nodes based on the node's betweenness, degree, or at random, and then calculating the percentage of remaining nodes (size) of the remaining connected network component. The fraction of nodes removed is shown on the x-axis and the fraction of remaining nodes is shown on the y-axis as the largest remaining network component size. Each color represents a depth zone while each line type represents the eukaryotic group assessed in each network. A more robust network is indicated by a larger area under the curve. The Protist curves overlap with All Taxa curves.

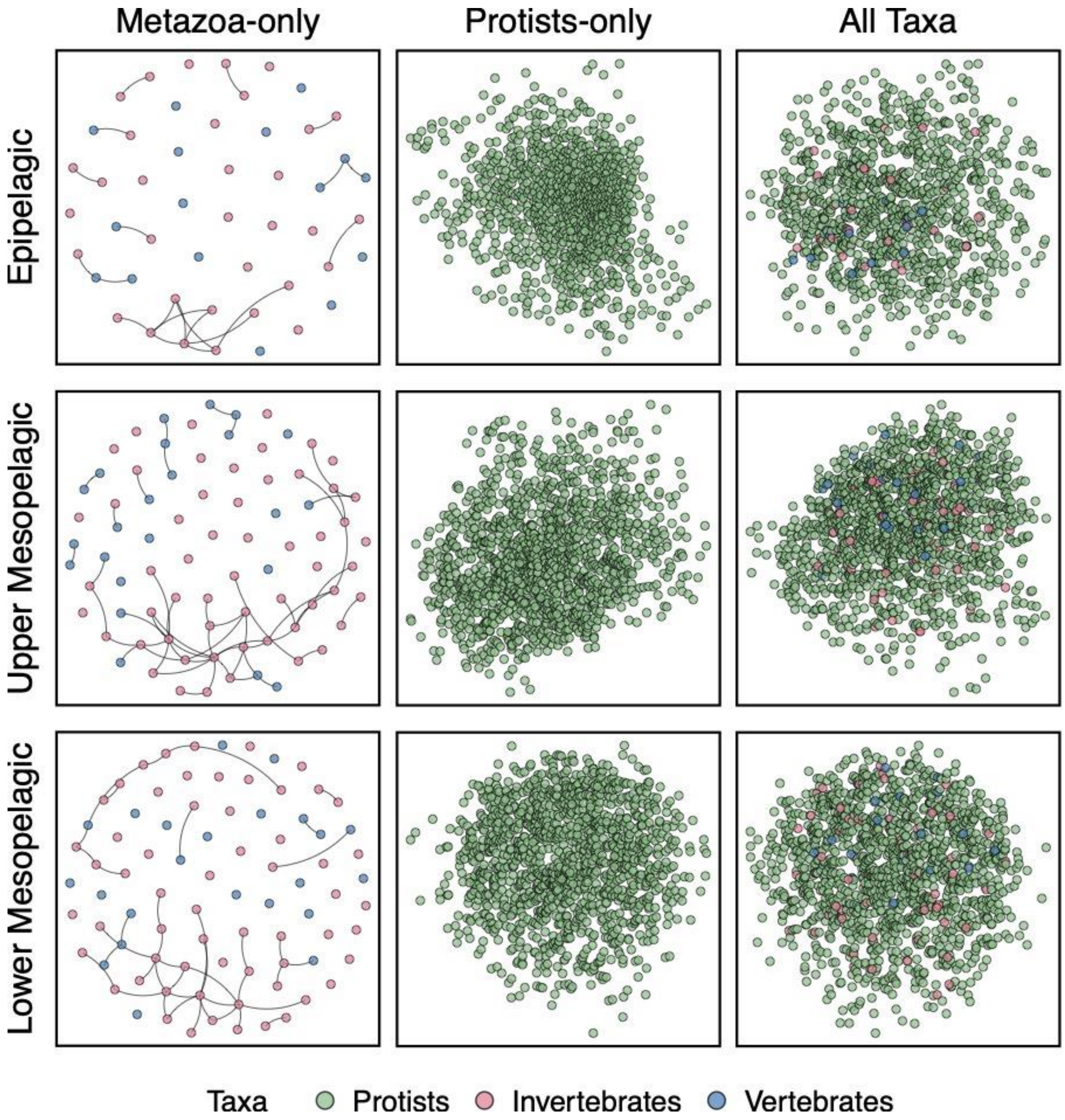


**Supplementary Figure S15**. A larger version of depth-specific networks (Figure 5) for Metazoa-only (invertebrate and vertebrates), Protists-only, and All Taxa (all ASVs). All nodes are shown as points with color indicating the broad taxonomic classification of protists, invertebrates, and vertebrates. Edges are shown for the Metazoa-only network to show the fragmented nature of the network with singleton nodes. The Protists-only and All Taxa networks are connected networks and edges are not shown. All networks include predicted positive and negative interactions.

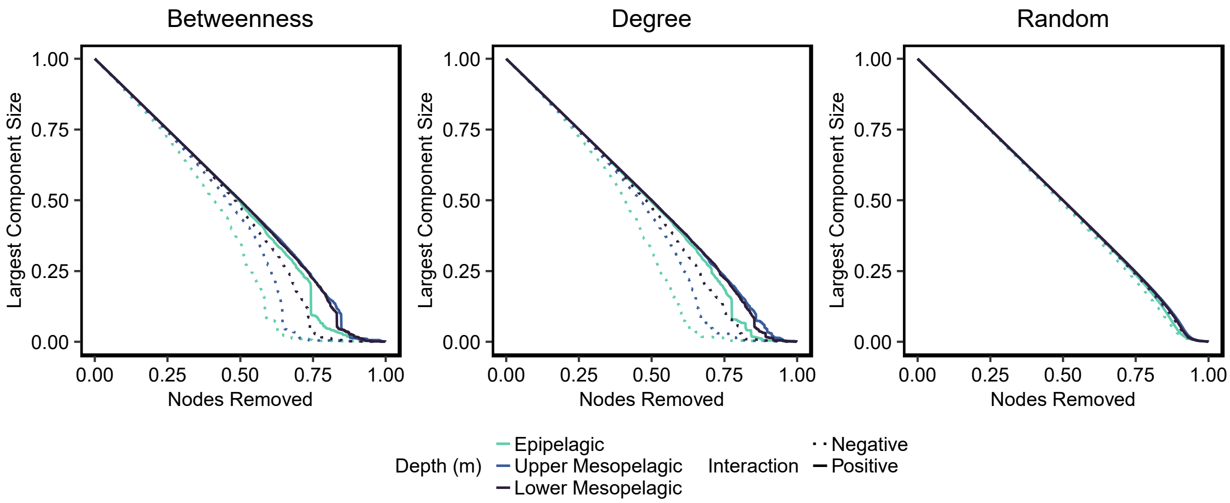


**Supplementary Figure S16.** Robustness curves at each depth zone for Negative and Positive networks parsed from the All Taxa network. Attack robustness of each network was measured by sequentially removing nodes based on the node's betweenness, degree, or at random, and then calculating the percentage of remaining nodes (size) of the remaining connected network component. The percentage of nodes removed is shown on the x-axis and the percentage of remaining nodes is shown on the y-axis as the largest remaining network component size. Each color represents a depth zone while each line type represents either the negative or positive network. A more robust network is indicated by a larger area under the curve.

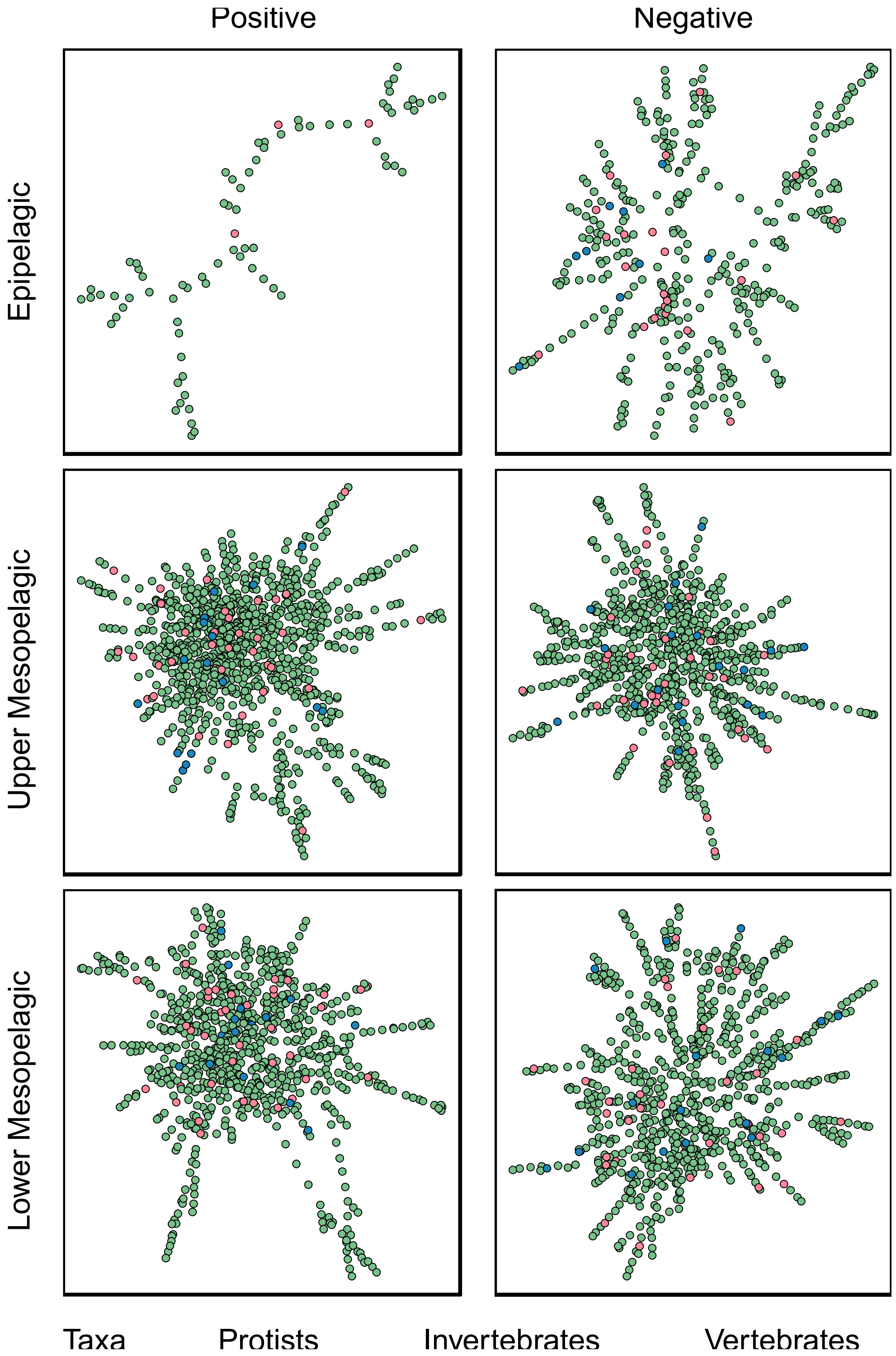


**Supplementary Figure S17.** Depth-specific All Taxa networks showing positive interactions (left) and negative interactions (right) comprised of nodes, with edge weights above the 90th percentile showing the strongest associations in each network. Each node represents an ASV, and the color of the nodes indicates the broad eukaryotic taxonomic group. Edges are shown as curved grey lines between nodes.

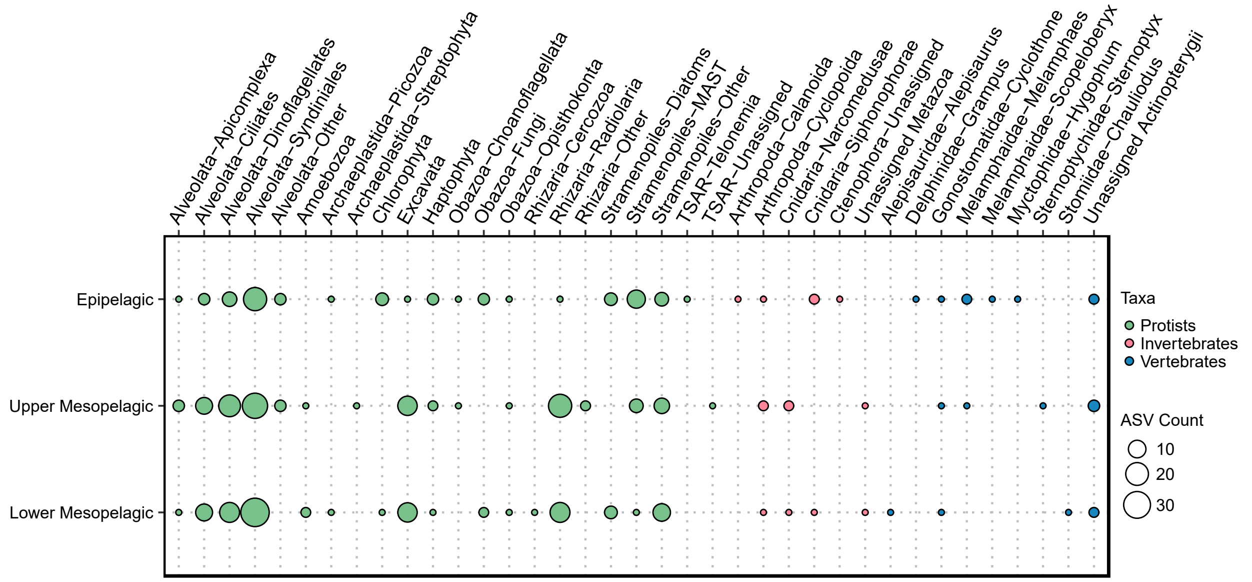


**Supplementary Figure S18.** ASVs above the 90th percentile in terms of degree and betweenness from the positive All Taxa network at each depth zone. The color of each point indicates the broader eukaryotic grouping while the size of the point indicates the number of ASVs that fall under each taxonomic subgroup. Taxonomic groups are listed on the x-axis and the depth is shown on the y-axis.

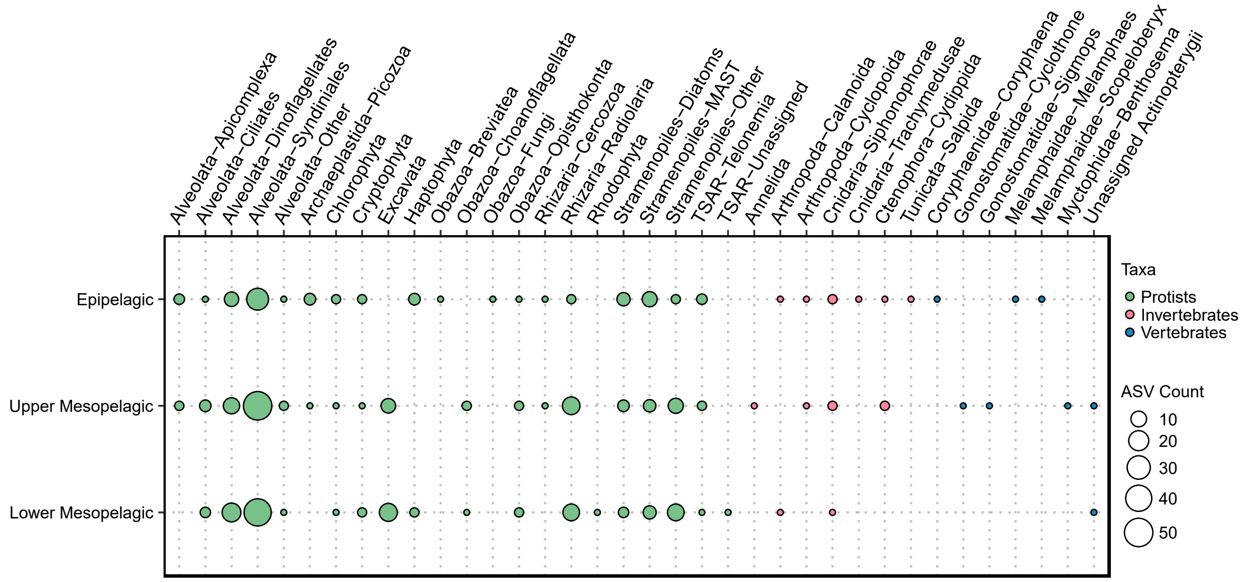


**Supplementary Figure S19.** ASVs above the 90th percentile in terms of degree and betweenness from the negative All Taxa network at each depth zone. The color of each point indicates the broader eukaryotic grouping while the size of the point indicates the number of ASVs that fall under each taxonomic subgroup. Taxonomic groups are listed on the x-axis and the depth is shown on the y-axis.

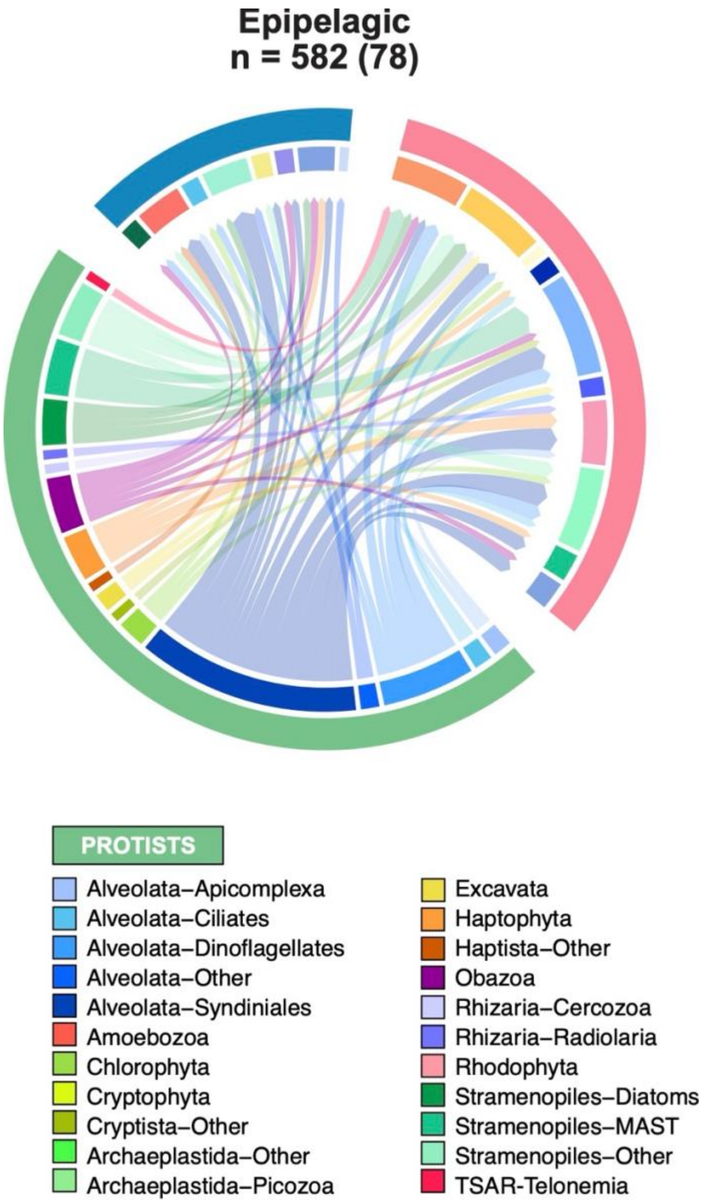


**Supplementary Figure S20.** A larger version of Figure 6. The epipelagic chord diagram indicates the number of strong pairwise interactions in the negative network. Interactions were selected by removing all ASV-ASV linkages below the 90$^{th}$ percentile threshold for edge weight (so that only the strongest interactions remained). Only protist-invertebrate, protist-vertebrate, and invertebrate-vertebrate interactions were considered. The total number of pairwise associations for each negative network is listed (n) with the number of strong associations in parentheses. Colors along the circumference (track) of the diagram indicate taxonomic groupings. The outer track shows protists in green, invertebrates in pink, and vertebrates in blue. The inner track further divides these groupings into taxonomic groups. Each link between two ASVs (or interaction) is depicted as a directional arrow that starts from the ASV in a lower trophic level and points towards the ASV at the higher trophic level. The arrows are also colored based on the lower trophic level ASV. Trophic levels are simplified and ordered as protist > invertebrate > vertebrate. The size of each inner track segment indicates the relative number of ASVs under each taxonomic group within a chord diagram.

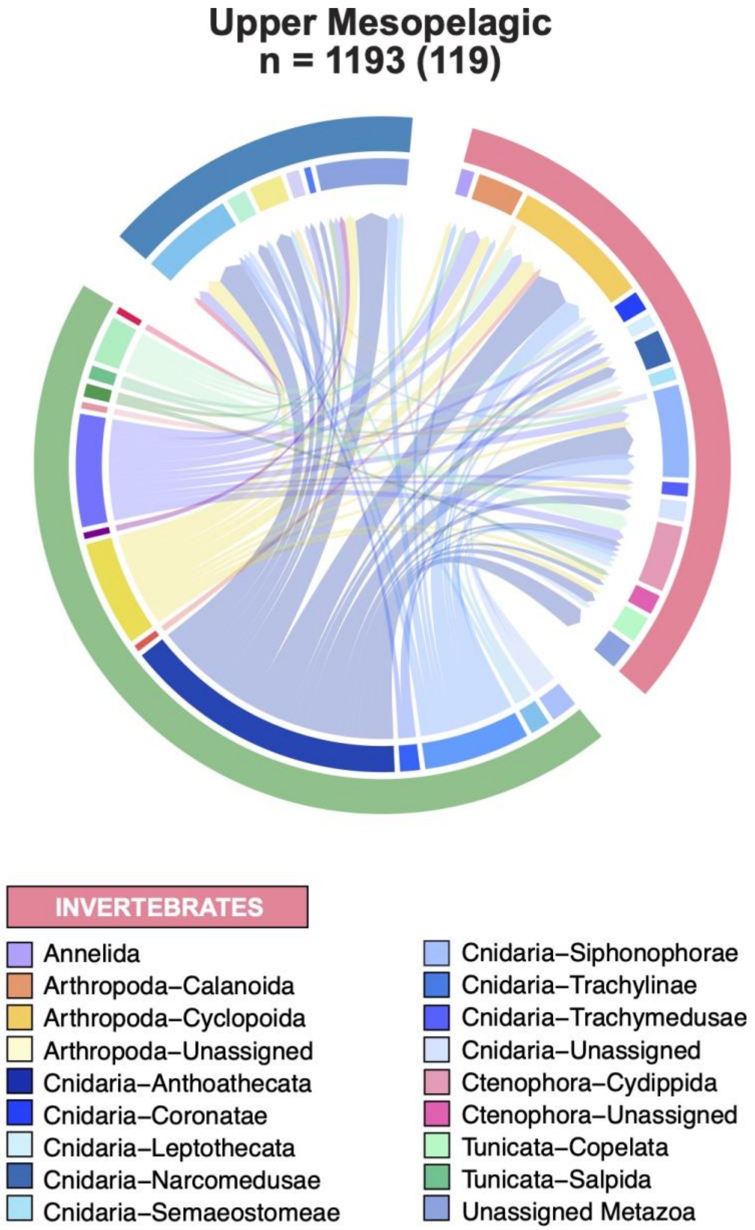


**Supplementary Figure S21.** A larger version of Figure 6. The upper mesopelagic chord diagram indicates the number of strong pairwise interactions in the negative network. Interactions were selected by removing all ASV-ASV linkages below the $90^{th}$ percentile threshold for edge weight (so that only the strongest interactions remained). Only protist-invertebrate, protist-vertebrate, and invertebrate-vertebrate interactions were considered. The total number of pairwise associations for each negative network is listed (n) with the number of strong associations in parentheses. Colors along the circumference (track) of the diagram indicate taxonomic groupings. The outer track shows protists in green, invertebrates in pink, and vertebrates in blue. The inner track further divides these groupings into taxonomic groups. Each link between two ASVs (or interaction) is depicted as a directional arrow that starts from the ASV in a lower trophic level and points towards the ASV at the higher trophic level. The arrows are also colored based on the lower trophic level ASV. Trophic levels are simplified and ordered as protist > invertebrate > vertebrate. The size of each inner track segment indicates the relative number of ASVs under each taxonomic group within a chord diagram.

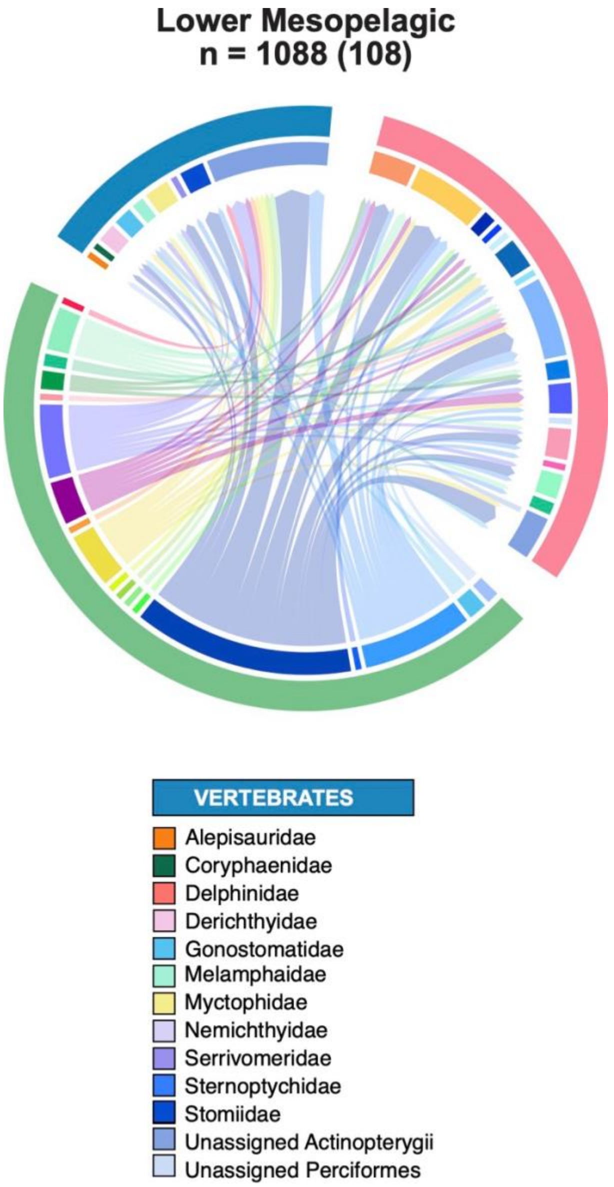


**Supplementary Figure S22.** A larger version of Figure 6. The lower mesopelagic chord diagram indicates the number of strong pairwise interactions in the negative network. Interactions were selected by removing all ASV-ASV linkages below the 90$^{th}$ percentile threshold for edge weight (so that only the strongest interactions remained). Only protist-invertebrate, protist-vertebrate, and invertebrate-vertebrate interactions were considered. The total number of pairwise associations for each negative network is listed (n) with the number of strong associations in parentheses. Colors along the circumference (track) of the diagram indicate taxonomic groupings. The outer track shows protists in green, invertebrates in pink, and vertebrates in blue. The inner track further divides these groupings into taxonomic groups. Each link between two ASVs (or interaction) is depicted as a directional arrow that starts from the ASV in a lower trophic level and points towards the ASV at the higher trophic level. The arrows are also colored based on the lower trophic level ASV. Trophic levels are simplified and ordered as protist > invertebrate > vertebrate. The size of each inner track segment indicates the relative number of ASVs under each taxonomic group within a chord diagram.

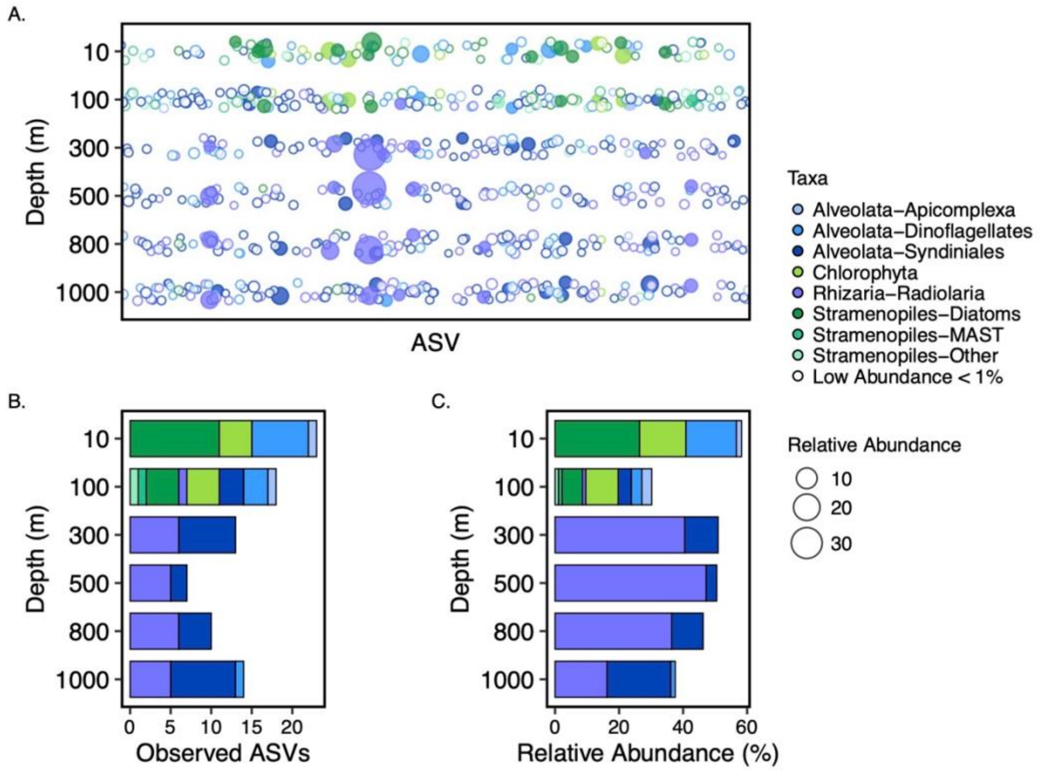


**Supplementary Figure S23**. A small portion of ASVs at each depth A) contribute a significant portion of the community relative abundance B). Within specific taxonomic groups, a select number of individual ASVs are highly abundant C). The colors indicate the protist taxonomic group. In A), the size of the point indicates relative abundance of an ASV. Points with no fill indicate low abundance ASVs and the outline color indicates taxa. ASVs at each depth with relative abundance > 1% are shown and all others are grouped under “Low Abundance”.

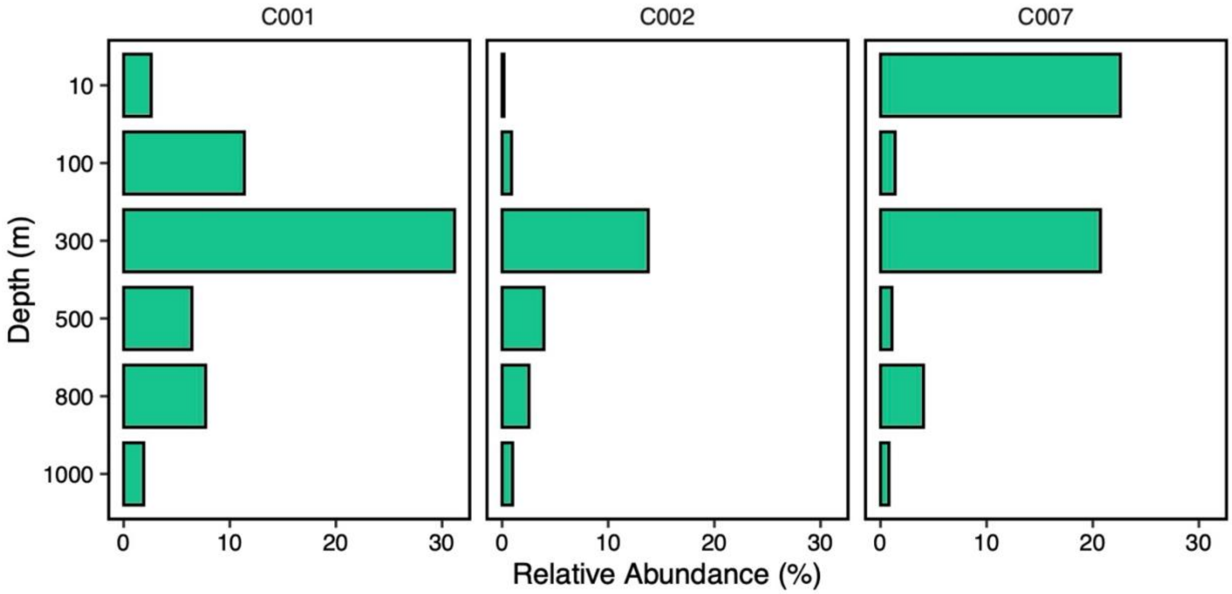


**Supplementary Figure S24.** Depth profiles of relative read abundance (%) for Tunicata-Salpida shown for each deployment. C001 and C007 were collected during the night and C002 was collected during the day.

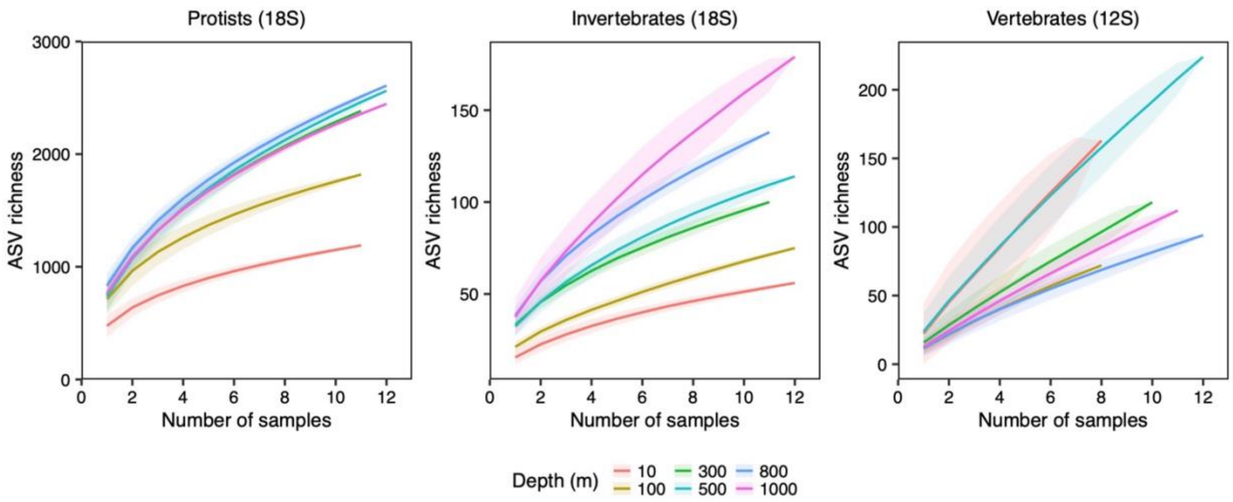


**Supplementary Figure S25.** Protist, invertebrate, and vertebrate ASV accumulation curves for each sampling depth. Solid lines represent the mean ASV richness calculated across 999 random permutations of sample order, and shaded areas indicate ±1 standard deviation across permutations, reflecting variability in richness accumulation with sampling effort.

### 5. Supplementary Tables

**Supplementary Table S1.** Coordinates (decimal degree) of sampling sites.

| CTD Cast | Latitude | Longitude |
|---|---|---|
| 001 | -70.88567 | 39.62350 |
| 002 | -70.88550 | 39.63583 |
| 007 | -70.96183 | 39.68050 |

**Supplementary Table S2.** The number of ASVs and relative read abundance for rare protistan taxa. Only taxa with < 1% relative read abundance are shown. ASVs were grouped into Supergroup and/or Division or Subdivision.

| Taxa | Observed ASVs | Relative Abundance (%) |
|---|---|---|
| Alveolata-Other | 221 | 0.960 |
| Amoebozoa | 29 | 0.140 |
| Archaeplastida-Other | 37 | 0.024 |
| Archaeplastida-Picozoa | 26 | 0.305 |
| Archaeplastida-Streptophyta | 32 | 0.010 |
| Cryptista-Kathablepharida | 6 | 0.133 |
| Cryptista-Other | 1 | 0.0001 |
| Cryptophyta | 25 | 0.783 |
| Eukaryota_X-Ancyromonadida | 1 | 0.0001 |
| Haptista-Centroplasthelida | 13 | 0.032 |
| Haptista-Other | 1 | 0.0001 |
| Obazoa-Apusomonada | 3 | 0.0002 |
| Obazoa-Breviatea | 2 | 0.005 |
| Obazoa-Choanoflagellata | 47 | 0.353 |
| Obazoa-Fungi | 136 | 0.310 |
| Obazoa-Opisthokonta | 148 | 0.402 |
| Provora-Nibbleridia | 1 | 0.00004 |
| Rhizaria-Cercozoa | 131 | 0.211 |
| Rhizaria-Other | 89 | 0.095 |
| Rhodophyta | 23 | 0.072 |
| TSAR-Other | 72 | 0.195 |
| TSAR-Telonemia | 56 | 0.270 |

**Supplementary Table S3.** The number of ASVs and relative abundance of invertebrate taxa with overall abundance < 1%. ASVs were grouped into Phylum-Order. Here, “Other” indicates unassigned Order.

| Taxa | Observed ASVs | Relative Abundance (%) |
|---|---|---|
| Annelida | 28 | 0.185 |
| Arthropoda-Amphipoda | 1 | 0.0005 |
| Arthropoda-Collembola | 1 | 0.0003 |
| Arthropoda-Decapoda | 1 | 0.0002 |
| Arthropoda-Eucarida | 2 | 0.0007 |
| Arthropoda-Halocyprida | 24 | 0.107 |
| Arthropoda-Harpacticoida | 3 | 0.0005 |
| Arthropoda-Mormonilloida | 1 | 0.139 |
| Arthropoda-Other | 17 | 0.054 |
| Arthropoda-Peracarida | 4 | 0.007 |
| Arthropoda-Sessilia | 2 | 0.002 |
| Bryozoa | 1 | 0.0001 |
| Chaetognatha | 5 | 0.004 |
| Cnidaria-Actiniaria | 1 | 0.0008 |
| Cnidaria-Anthoathecata | 9 | 0.154 |
| Cnidaria-Ceriantharia | 2 | 0.0003 |
| Cnidaria-Coronatae | 2 | 0.353 |
| Cnidaria-Leptothecata | 6 | 0.031 |
| Cnidaria-Narcomedusae | 12 | 0.230 |
| Cnidaria-Other | 9 | 0.126 |
| Cnidaria-Semaeostomeae | 7 | 0.164 |
| Cnidaria-Trachylinae | 6 | 0.089 |
| Cnidaria-Trachymedusae | 7 | 0.386 |
| Ctenophora-Other | 14 | 0.121 |
| Ctenophora-Platyctenida | 1 | 0.001 |
| Echinodermata | 4 | 0.002 |
| Gastrotricha | 1 | 0.015 |
| Hemichordata | 1 | 0.0004 |
| Mollusca | 7 | 0.024 |
| Nematoda | 2 | 0.0009 |
| Nemertea | 1 | 0.002 |
| Platyhelminthes | 1 | 0.0002 |
| Porifera | 1 | 0.0001 |
| Rotifera | 8 | 0.003 |
| Tunicata-Doliolida | 2 | 0.0005 |
| Unassigned Metazoa | 59 | 0.300 |

**Supplementary Table S4.** The number of ASVs and relative abundance of vertebrate taxa with overall abundance < 1%. ASVs were grouped into Family.

| Taxa | Observed ASVs | Relative Abundance (%) |
|---|---|---|
| Alepisauridae | 2 | 0.047 |
| Ammodytidae | 3 | 0.034 |
| Balaenopteridae | 2 | 0.005 |
| Bathylagidae | 3 | 0.267 |
| Caristiidae | 1 | 0.026 |
| Coryphaenidae | 5 | 0.124 |
| Delphinidae | 3 | 0.033 |
| Derichthyidae | 2 | 0.029 |
| Eurypharyngidae | 1 | 0.015 |
| Gempylidae | 2 | 0.258 |
| Howellidae | 3 | 0.020 |
| Microstomatidae | 3 | 0.130 |
| Moridae | 1 | 0.683 |
| Phosichthyidae | 2 | 0.256 |
| Phycidae | 1 | 0.0002 |
| Scombridae | 2 | 0.474 |
| Stromateidae | 1 | 0.142 |
| Unassigned Actinopterygii | 222 | 0.945 |
| Unassigned Perciformes | 6 | 0.004 |
| Zoarcidae | 3 | 0.609 |

**Supplementary Table S5.** Analysis of Variance test results for alpha diversity indices (stats) across depths. The indices included in the test were Richness (observed ASVs), Shannon Index (Shannon) and Pielou's Evenness Index (Evenness). NA is *Not Applicable.*

| taxa | Stats | term | df | sumsq | meansq | statistic | p.value |
|---|---|---|---|---|---|---|---|
| **Protists** | Richness | depth | 5 | 821962.5 | 164392.5 | 12.7 | 1.57E-08 |
| | | Residuals | 63 | 816889.7 | 12966.5 | NA | NA |
| | Shannon | depth | 5 | 15.7 | 3.1 | 65.6 | 1.16E-23 |
| | | Residuals | 63 | 3.0 | 0.0 | NA | NA |
| | Evenness | depth | 5 | 0.4 | 0.1 | 112.7 | 4.57E-30 |
| | | Residuals | 63 | 0.0 | 0.0 | NA | NA |
| **Invertebrates** | Richness | depth | 5 | 5177.3 | 1035.5 | 31.6 | 4.62E-16 |
| | | Residuals | 64 | 2098.1 | 32.8 | NA | NA |
| | Shannon | depth | 5 | 16.4 | 3.3 | 17.8 | 5.36E-11 |
| | | Residuals | 64 | 11.8 | 0.2 | NA | NA |
| | Evenness | depth | 5 | 0.8 | 0.2 | 10.4 | 2.35E-07 |
| | | Residuals | 64 | 1.0 | 0.0 | NA | NA |
| **Vertebrates** | Richness | depth | 5 | 1328.8 | 265.8 | 2.2 | 0.07 |
| | | Residuals | 55 | 6725.0 | 122.3 | NA | NA |
| | Shannon | depth | 5 | 4.4 | 0.9 | 1.6 | 0.18 |
| | | Residuals | 55 | 30.3 | 0.6 | NA | NA |
| | Evenness | depth | 5 | 0.4 | 0.1 | 1.4 | 0.23 |
| | | Residuals | 55 | 3.0 | 0.1 | NA | NA |

**Supplementary Table S6**. Tukey's Honestly Significant Difference (HSD) test results for protist alpha diversity indices (stats) between depths (contrast) following the ANOVA test (Supplementary Table S1). The indices included in the test were Richness (observed ASVs), Shannon Index (Shannon) and Pielou's Evenness Index (Evenness).

| taxa | stats | contrast | estimate | conf.low | conf.high | p.value |
|---|---|---|---|---|---|---|
| | | 100-10 | 242.7 | 100.0 | 385.4 | 6.96E-05 |
| | | 500-10 | 261.9 | 122.2 | 401.6 | 1.03E-05 |
| | Richness | 300-10 | 277.5 | 134.8 | 420.2 | 4.70E-06 |
| | | 1000-10 | 293.2 | 153.5 | 432.9 | 8.07E-07 |
| | | 800-10 | 345.3 | 205.6 | 485.0 | 1.03E-08 |
| | | 300-500 | 0.3 | 0.0 | 0.5 | 3.86E-02 |
| | | 10-500 | 0.4 | 0.2 | 0.7 | 2.69E-04 |
| | | 800-500 | 0.5 | 0.3 | 0.8 | 9.43E-07 |
| | | 1000-500 | 1.2 | 0.9 | 1.4 | 1.53E-11 |
| | | 100-500 | 1.4 | 1.1 | 1.6 | 1.53E-11 |
| | Shannon | 800-300 | 0.3 | 0.0 | 0.5 | 4.82E-02 |
| | | 1000-300 | 0.9 | 0.6 | 1.1 | 1.64E-11 |
| | | 100-300 | 1.1 | 0.8 | 1.3 | 1.53E-11 |
| | | 1000-10 | 0.7 | 0.5 | 1.0 | 5.67E-10 |
| | | 100-10 | 0.9 | 0.7 | 1.2 | 1.56E-11 |
| **Protists** | | 1000-800 | 0.6 | 0.3 | 0.9 | 7.43E-08 |
| | | 100-800 | 0.8 | 0.5 | 1.1 | 3.51E-11 |
| | | 300-500 | 0.0 | 0.0 | 0.1 | 4.05E-03 |
| | | 800-500 | 0.1 | 0.0 | 0.1 | 4.79E-08 |
| | | 10-500 | 0.1 | 0.1 | 0.1 | 1.54E-11 |
| | | 1000-500 | 0.2 | 0.1 | 0.2 | 1.53E-11 |
| | | 100-500 | 0.2 | 0.2 | 0.2 | 1.53E-11 |
| | | 10-300 | 0.1 | 0.0 | 0.1 | 4.12E-08 |
| | Evenness | 1000-300 | 0.1 | 0.1 | 0.2 | 1.53E-11 |
| | | 100-300 | 0.2 | 0.1 | 0.2 | 1.53E-11 |
| | | 10-800 | 0.0 | 0.0 | 0.1 | 1.35E-03 |
| | | 1000-800 | 0.1 | 0.1 | 0.1 | 1.65E-11 |
| | | 100-800 | 0.1 | 0.1 | 0.2 | 1.53E-11 |
| | | 1000-10 | 0.1 | 0.0 | 0.1 | 3.68E-05 |
| | | 100-10 | 0.1 | 0.1 | 0.1 | 4.40E-11 |
| | | 100-1000 | 0.0 | 0.0 | 0.1 | 5.07E-03 |

**Supplementary Table S7**. Tukey's Honestly Significant Difference (HSD) test results for invertebrate alpha diversity indices (stats) between depths (contrast) following the ANOVA test (Supplementary Table S1). The indices included in the test were Richness (observed ASVs), Shannon Index (Shannon) and Pielou’s Evenness Index (Evenness). Note: indices were not significant for vertebrates and are not included.

| taxa | stats | contrast | estimate | conf.low | conf.high | p.value |
|---|---|---|---|---|---|---|
| Invertebrates | Richness | 500-10 | 17.0 | 10.1 | 23.9 | 9.28E-09 |
| | | 300-10 | 18.7 | 11.7 | 25.8 | 9.27E-10 |
| | | 1000-10 | 21.6 | 14.8 | 28.5 | 1.26E-11 |
| | | 800-10 | 24.0 | 16.9 | 31.0 | 9.73E-12 |
| | | 500-100 | 10.6 | 3.7 | 17.5 | 3.65E-04 |
| | | 300-100 | 12.3 | 5.3 | 19.4 | 3.70E-05 |
| | | 1000-100 | 15.2 | 8.4 | 22.1 | 1.93E-07 |
| | | 800-100 | 17.6 | 10.5 | 24.6 | 6.79E-09 |
| | Shannon | 100-10 | 0.6 | 0.1 | 1.1 | 8.64E-03 |
| | | 1000-10 | 0.6 | 0.1 | 1.2 | 7.24E-03 |
| | | 500-10 | 1.1 | 0.5 | 1.6 | 1.17E-06 |
| | | 300-10 | 1.2 | 0.7 | 1.7 | 9.44E-08 |
| | | 800-10 | 1.5 | 1.0 | 2.0 | 1.09E-10 |
| | Evenness | 300-100 | 0.6 | 0.0 | 1.1 | 2.46E-02 |
| | | 800-100 | 0.9 | 0.4 | 1.4 | 9.73E-05 |
| | | 300-1000 | 0.6 | 0.0 | 1.1 | 2.88E-02 |

**Supplementary Table S8.** Network topology for depth- and tax-specific networks. All Taxa networks integrate protist, invertebrate, and vertebrate ASVs. Protist networks only contain protist ASVs. Metazoa networks integrate invertebrates and vertebrates. Edge density is calculated as the edge-to-node ratio. N.conn = node connectivity and E.conn = edge connectivity.

| Depth | Taxa | Nodes | Edges | Edge Density | N.conn | E.conn |
|---|---|---|---|---|---|---|
| **Epipelagic** | All | 1040 | 12279 | 11.8 | 3 | 3 |
| | Protist | 991 | 11245 | 11.3 | 4 | 4 |
| | Metazoa | 49 | 21 | 0.43 | 0 | 0 |
| **Upper Mesopelagic** | All | 1424 | 23338 | 16.4 | 4 | 4 |
| | Protist | 1345 | 21274 | 15.8 | 4 | 4 |
| | Metazoa | 79 | 53 | 0.67 | 0 | 0 |
| **Lower Mesopelagic** | All | 1397 | 21807 | 15.6 | 13 | 13 |
| | Protist | 1314 | 19619 | 14.9 | 10 | 10 |
| | Metazoa | 83 | 41 | 0.49 | 0 | 0 |

**Supplementary Table S9**. Network topology for depth-specific networks integrating all taxonomic groups and parsed out by interaction type. “Both” networks include both positive and negative interactions. Edge % is calculated as the proportion of positive or negative edges relative to all edges in the network. Edge density is calculated as the edge-to-node ratio.

| Depth | Interaction Type | Nodes | Edges | Edge % | Edge Density | Modularity |
|---|---|---|---|---|---|---|
| **Epipelagic** | Both | 1040 | 12279 | | 11.8 | 0.26 |
| | Positive | 1040 | 7117 | 58% | 6.8 | 0.41 |
| | Negative | 1016 | 5162 | 42% | 5.1 | 0.29 |
| **Upper Mesopelagic** | Both | 1424 | 23338 | | 16.4 | 0.21 |
| | Positive | 1424 | 13301 | 57% | 9.3 | 0.33 |
| | Negative | 1423 | 10037 | 43% | 7.1 | 0.21 |
| **Lower Mesopelagic** | Both | 1397 | 21807 | | 15.6 | 0.18 |
| | Positive | 1397 | 11611 | 53% | 8.3 | 0.31 |
| | Negative | 1397 | 10196 | 47% | 7.3 | 0.19 |

**Supplementary Table S10.** Node connectivity (N.conn) and edge connectivity (E.conn) for networks including positive and negative interactions (Both), positive interactions only (Positive), and negative interactions only (Negative).

| Depth | Interaction Type | N.conn | E.conn |
|---|---|---|---|
| **Epipelagic** | Both | 3 | 3 |
| | Positive | 3 | 3 |
| | Negative | 1 | 1 |
| **Upper Mesopelagic** | Both | 4 | 4 |
| | Positive | 3 | 3 |
| | Negative | 1 | 1 |
| **Lower Mesopelagic** | Both | 13 | 13 |
| | Positive | 2 | 2 |
| | Negative | 2 | 2 |

**Supplementary Table S11.** Vertebrate species detected by the 12S MiFish primer across three CTD casts and 6 depths. Deep-sea fish species found in the surface (< 200 m) are indicated with an asterisk*.

| Depth | Cast 1 | Cast 2 | Cast 7 |
|---|---|---|---|
| **10 m** | *Mola mola* | *Melamphaes microps** | *Serrivomer beanii** |
| | *Hygophum proximum** | *Scopeloberyx opisthopterus** | *Thunnus thynnus* |
| | *Delphinus delphis* | *Leptostomias robustus** | *Melamphaes microps** |
| | *Grampus griseus* | *Sternoptyx diaphana** | *Stomias boa** |
| | | *Sternoptyx pseudodiaphana** | *Benthosema glaciale** |
| | | *Hygophum proximum** | *Scopeloberyx opisthopterus** |
| | | | *Leptostomias robustus** |
| | | | *Sternoptyx diaphana** |
| **100 m** | *Bajacalifornia megalops** | *Benthosema glaciale** | *Serrivomer beanii** |
| | *Cyclothone pallida** | *Cyclothone microdon** | *Melamphaes microps** |
| | *Melamphaes microps** | *Melamphaes microps** | *Cyclothone microdon** |
| | *Cyclothone microdon** | *Leptostomias robustus** | *Coryphaena hippurus* |
| | *Coryphaena hippurus* | *Scopeloberyx opisthopterus** | *Scopeloberyx opisthopterus** |
| | | *Balaenoptera acutorostrata* | |
| **300 m** | *Notoscopelus kroyeri* | *Benthosema glaciale* | *Sigmops elongatus* |
| | *Cyclothone microdon* | *Cyclothone microdon* | *Lampadena speculigera* |
| | *Melamphaes microps* | *Sigmops elongatus* | *Polyipnus polli* |
| | *Scopelogadus mizolepis* | *Valenciennellus tripunctulatus* | *Notoscopelus kroyeri* |
| | *Stenella coeruleoalba* | *Argyropelecus aculeatus* | *Scopeloberyx opisthopterus* |
| | | *Scopelogadus beanii* | *Cyclothone microdon* |
| | | | *Melamphaes microps* |
| | | | *Leptostomias robustus* |
| | | | *Sternoptyx pseudodiaphana* |
| | | | *Howella brodiei* |
| | | | *Sternoptyx diaphana* |
| | | | *Hygophum hygomii* |
| **500 m** | *Cyclothone braueri* | *Benthosema glaciale* | *Cyclothone microdon* |
| | *Lampadena speculigera* | *Diaphus metopoclampus* | *Cyclothone braueri* |
| | *Benthosema glaciale* | *Serrivomer beanii* | *Melamphaes microps* |
| | *Melamphaes microps* | *Sigmops elongatus* | *Lampadena speculigera* |
| | *Cyclothone microdon* | *Cyclothone braueri* | *Sigmops elongatus* |
| | *Ichthyococcus ovatus* | *Nansenia ardesiaca* | *Benthosema glaciale* |
| | *Scopeloberyx opisthopterus* | *Scopelogadus beanii* | *Leptostomias robustus* |
| | *Leptostomias robustus** | *Chauliodus sloani* | *Scopeloberyx opisthopterus* |

| | | | |
|---|---|---|---|
| | *Myctophum affine* | *Coryphaena hippurus* | *Sternoptyx diaphana* |
| | *Cyclothone pseudopallida* | *Benthosema suborbitale* | *Melamphaes lugubris* |
| | *Notoscopelus kroyeri* | *Nemichthys curvirostris* | *Scopelogadus mizolepis* |
| | *Scopelogadus beanie* | *Symbolophorus veranyi* | *Howella brodiei* |
| | *Diaphus gigas* | *Ceratoscopelus maderensis* | *Nannobrachium lineatum* |
| | *Sternoptyx diaphana* | *Melamphaes microps* | *Ammodytes dubius* |
| | *Nemichthys curvirostris* | | *Bolinichthys pyrsobolus* |
| | *Chauliodus sloani* | | *Sternoptyx pseudodiaphana* |
| | *Ceratoscopelus maderensis* | | *Chauliodus sloani* |
| | | | *Ceratoscopelus maderensis* |
| | | | *Argyropelecus hemigymnus* |
| **800 m** | *Benthosema glaciale* | *Sigmops elongatus* | *Cyclothone microdon* |
| | *Sigmops elongatus* | *Cyclothone microdon* | *Benthosema glaciale* |
| | *Melanostigma atlanticum* | *Benthosema glaciale* | *Serrivomer beanii* |
| | *Melamphaes microps* | *Scopeloberyx opisthopterus* | *Scopeloberyx opisthopterus* |
| | *Cyclothone microdon* | *Serrivomer beanii* | *Lampadena speculigera* |
| | *Lipolagus ochotensis* | *Chauliodus sloani* | *Scopelogadus mizolepis* |
| | *Nansenia ardesiaca* | *Alepisaurus ferox* | *Melamphaes microps* |
| | *Serrivomer beanii* | *Ammodytes dubius* | *Ammodytes dubius* |
| | *Scopeloberyx opisthopterus* | *Nessorhamphus ingolfianus* | *Nessorhamphus ingolfianus* |
| | *Poromitra capito* | *Lampanyctus macdonaldi* | *Nannobrachium cuprarium* |
| | *Lampanyctus macdonaldi* | | *Balaenoptera physalus* |
| | *Urophycis tenuis* | | *Derichthys serpentinus* |
| **1000 m** | *Scopeloberyx opisthopterus* | *Benthosema glaciale* | *Serrivomer beanii* |
| | *Laemonema longipes* | *Leptostomias robustus* | *Notoscopelus resplendens1* |
| | *Photostomias guernei* | *Scopeloberyx opisthopterus* | *Cyclothone microdon* |
| | *Benthosema glaciale* | *Melamphaes microps* | *Melamphaes microps* |
| | *Nealotus tripes* | *Eurypharynx pelecanoides* | *Leptostomias robustus* |
| | *Cyclothone microdon* | *Astronesthes_X* | *Scopeloberyx opisthopterus* |
| | *Lipolagus ochotensis* | *Nessorhamphus ingolfianus* | *Benthosema glaciale* |
| | *Leptostomias robustus* | *Sternoptyx pseudodiaphana* | *Chauliodus sloani* |
| | *Melamphaes microps* | *Sternoptyx diaphana* | *Coryphaena hippurus* |
| | *Chauliodus sloani* | *Cyclothone microdon* | *Alepisaurus ferox* |
| | *Sternoptyx pseudodiaphana* | *Nemichthys curvirostris* | |
| | *Bathylagus pacificus* | | |
| | *Sternoptyx diaphana* | | |
| | *Coryphaena hippurus* | | |
| | *Sigmops elongatus* | | |
